# Meta-learning is expressed through altered prefrontal cortical dynamics

**DOI:** 10.64898/2026.01.01.697272

**Authors:** Xulu Sun, Alison E. Comrie, Ari E. Kahn, Emily J. Monroe, Clayton B. Washington, Abhilasha Joshi, Jennifer A. Guidera, Eric L. Denovellis, Timothy A. Krausz, Jenny Zhou, Paige Thompson, Jose Hernandez, Allison Yorita, Razi Haque, Chethan Pandarinath, Joshua D. Berke, Nathaniel D. Daw, Loren M. Frank

**Affiliations:** Department of Physiology and Psychiatry, University of California, San Francisco; San Francisco, CA 94158, USA; Howard Hughes Medical Institute; Chevy Chase, MD 20815, USA; Neuroscience Graduate Program, University of California San Francisco; San Francisco, CA 94158, USA; Princeton Neuroscience Institute, Princeton University; Princeton, NJ 08544, USA; Center for Machine Learning and Department of Biomedical Engineering, Georgia Institute of Technology and Emory University; Atlanta, GA 30322, USA; Medical Scientist Training Program, University of California, San Francisco; San Francisco, CA 94143, USA; Center for Micro- and Nano-Technology, Lawrence Livermore National Laboratory; Livermore, 94550, USA; Kavli Institute for Fundamental Neuroscience, University of California, San Francisco; San Francisco, CA 94158, USA; Department of Neurology and Department of Psychiatry and Behavioral Science, and Weill Institute for Neurosciences, University of California, San Francisco; San Francisco, CA 94158, USA; Department of Psychology, Princeton University; Princeton, NJ 08544, USA

## Abstract

Learning where and when rewards like food and water are available is essential for survival^1,2^. In the simplest cases where resource availability is stable, animals can learn reward contingencies by integrating outcomes across repeated samples of each possible action. In more natural settings, however, reward availability is governed by structured higher-order rules such as depletion and repletion over time. To adapt flexibly to such changing environments, optimal choices require meta-learning wherein animals learn how to learn from external feedback, ultimately enabling them to infer the underlying reward structure from abstract, generalizable rules rather than relying solely on recent outcomes^3,4^. The existence of meta-learning in animal behavior is well established^3–8^, yet the neural circuits and computations that implement it remain poorly understood^9–11^. Here we investigated meta-learning using a spatial foraging task in which rats acquired a depletion-repletion rule that regulated reward availability, and carried out longitudinal, high-density recordings from the medial prefrontal cortex (mPFC). We show that meta-learning engages specific, systematic changes in mPFC neural dynamics that embed the learned rule and thereby alter how the network learns action values from reward outcomes. These dynamics are based on mixed coding of task structure and value in individual mPFC neurons. At the population level, this coding organizes into low-dimensional dynamical motifs that generalize across task conditions. As meta-learning progresses, these motifs are reshaped to instantiate both rule-guided inference of future states before outcome delivery and rule-based value updating during the outcome period. These results indicate that meta-learning sculpts pre-existing prefrontal dynamics to support the acquisition of new, generalizable reward-learning strategies.

## Main

To make adaptive decisions in a changing environment, the brain must learn to incorporate external feedback into its evaluation processes and action policies. For instance, effective foraging in the wilderness requires animals to learn how food, water, or other rewards are distributed and use this knowledge to predict the likely outcomes of different choices^1,2^. In stable environments, such predictions are straightforward: repeated sampling of each possible action eventually converges on accurate estimates of reward probabilities. However, many natural environments impose additional structure—such as resource depletion with use and recovery over time—that demands learning not only immediate reward availability but also the rules governing how that availability evolves. This higher-order learning process, known as meta-learning, enables the brain to improve its learning strategies by abstracting regularities across experiences^3,4,12,13^. Through meta-learning, animals can refine predictions of action values and consequently make more advantageous choices. The resulting strategies can transfer to other contexts that share common rules^11,14,15^.

Converging evidence from recent experimental and theoretical studies points to the prefrontal cortex (PFC) as a critical substrate for meta-learning. Empirical work shows that PFC is necessary for refining learning algorithms and stabilizing value coding over experience^10^. Complementary computational models of prefrontal recurrent networks demonstrate that these circuits can implement rapid value updating, while the mechanism of this updating process—effectively the learning algorithm itself—is shaped by slower meta-learning that incorporates higher-order task statistics^9,16^. Such statistics often define how latent task states evolve over time, enabling rule-based prediction beyond immediate outcomes^9,17^. Further, PFC has been implicated in abstraction and generalization^18–20^, consistent with the proposition that meta-learned strategies could be expressed flexibly across different task conditions or contexts^14,21^. Despite these advances, it remains unclear whether meta-learning drives systematic changes in PFC population dynamics to implement altered value computation and support rule-based inference of future task states. Whether similar dynamics are reused to subserve generalized computations across contexts is also unknown.

Here we focus on the medial PFC (mPFC), a subregion known to represent abstract task structure, encode action-outcome contingencies, and predict future states^2,13,18,19,22–25^. We test the hypothesis that meta-learning sculpts mPFC population dynamics, altering the computations that underlie task-state inference, value updating, and action selection. We specifically sought dynamical mechanisms capable of flexibly implementing new learning rules. These potential mechanisms extend classic views of reward learning in the brain, wherein error-driven synaptic plasticity (e.g., at corticostriatal connections^26^) carries out reinforcement learning essentially through recency-weighted value computation^27^. Despite much empirical support for such classic learning rules, it remains unresolved how they can be adapted to situations where an action’s current value is dissociated from its recent outcomes.

We used a multi-patch foraging task in which rats first learned stable reward probabilities for each patch primarily by caching reward history^28^. In a later phase of the task, they acquired a depletion-repletion rule that structured reward availability across patches and dissociated reward expectancy from recent experience. We developed a statistical behavioral model to infer trial-by-trial subjective values predictive of rat decisions and to quantify the meta-learning process underlying shifts in reward-learning strategy. This task and model, combined with stable, high-density polymer probe recordings from mPFC over many tens of sessions^29^, enabled us to capture the reorganization of mPFC population dynamics as animals adapted their learning strategy to incorporate the rule.

We found that mPFC neurons simultaneously tracked task structure (e.g., goal progress, action, and foraging patch identity) and value. Building upon the single-neuron mixed-coding profiles, coordinated population activity formed dynamical motifs that instantiated value- and decision-relevant computations and were reused across task conditions. Crucially, meta-learning reshaped these motifs to support rule-based value inference, paralleling the emergence of a new reward-learning strategy. These results point to a general computational principle whereby structured population dynamics internalize task rules and flexibly adapt reward-learning algorithms across contexts. More broadly, the identification of recurring dynamical motifs illuminates how generalizable computations can be implemented across experiences, a core property of intelligence^30–32^.

### Behavioral evidence for meta-learning in the spatial bandit task

We developed a variant of a spatial bandit task^28^ in which rats (n = 4) foraged for probabilistic rewards distributed across multiple patches (Fig. 1a; see Methods). The environment consisted of three Y-shaped ‘foraging patches’ radiating from the maze center, and each patch contained two reward ports at path ends. The multiple bifurcations and reward sites exposed animals to a broader set of decision-making conditions than standard two-arm bandit task designs. This richer structure enables identification of generalized computational principles proposed in recent theoretical work^30,31^.

**Fig. 1.**
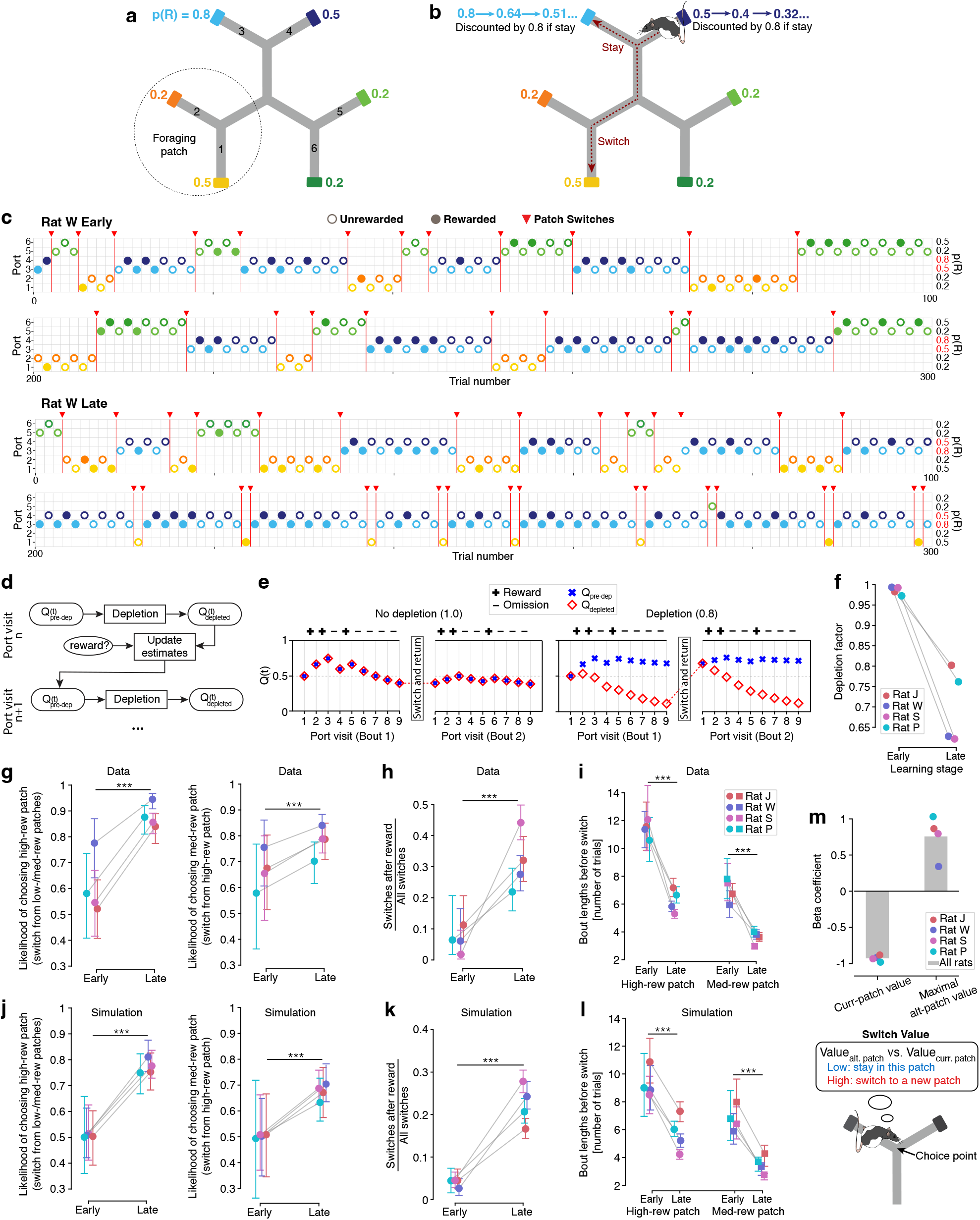
Rats expressed value-guided behavior in a spatial bandit task and learned a novel strategy. **a**, Maze schematic. Three Y-shaped foraging patches radiate from a central junction, and each patch contains two reward ports at path ends. Ports deliver liquid rewards according to assigned reward probabilities [p(R)] that vary across sessions. An example set of p(R) is shown. Ports are labeled as 1-6 and colored by their identity for figure visualization only. **b**, Task rules. Consecutive visits within the same patch (‘Stay’) discount p(R) by a factor of 0.8 after each visit. Visiting a different patch (‘Switch’) restores p(R) to its nominal value. Example depletion is illustrated for staying in the blue patch. **c**, Example behavior from the first and last 100 trials of representative early- and late-learning sessions. Filled circles: rewarded trials; hollow circles: unrewarded trials (colored by port identity); red triangles: patch switches. **d**, Schematic of value updating in the behavioral model with depletion. Subjective values of a visited port were first estimated as the posterior mean of a Beta distribution updated by reward outcomes (baseline value Q_pre-dep_). A depletion factor then discounted Q_pre-dep_ during repeated visits, yielding value estimates of the current trial (Q_depleted_) that reset upon patch switching. **e**, Example Q value estimation for a given port with and without the depletion factor. Red diamonds indicate the subjective value estimates on the current trial; blue crosses indicate belief about the baseline (undepleted) value; plus/minus signs indicate reward/omission. Left: without incorporating the effect of depletion, the baseline value is underestimated. Right: by quantifying the effect of depletion, the baseline value remains near its true undepleted level due to reduced influence of omissions, while trial-by-trial value estimates decrease across consecutive visits. *For illustration, a patch is simplified as one port, while in the real task, each patch consists of two ports and Q*_*patch*_ *is a weighted average of its two ports*. **f**, Depletion factor inferred from behavior decreased across days, indicating acquisition of the depletion-repletion rule. **g-i**, Behavioral metrics distinguishing early and late learning reflect the meta-learned new strategy. Error bars: 95% CI. **g**, Left: Increased likelihood of choosing the high-reward patch when switching from other patches (two-proportion Z-test per animal: P_J_ = 1.94 × 10^-7^, P_W_ = 5.10 × 10^-5^, P_S_ = 3.81 × 10^-8^, P_P_ = 5.30 × 10^-5^; linear mixed effects model across all animals: P = 1.99 × 10^-19^). Right: Increased likelihood of choosing the medium-reward patch when switching from the high-reward patch (two-proportion Z-test per animal: P_J_ = 0.08, P_W_ = 0.09, P_S_ = 0.05, P_P_ = 0.14; linear mixed effects model across animals: P = 1.93 × 10^-35^). **h**, Increased probability of switching after reward (two-proportion Z-test per animal: P_J_ = 0.00044, P_W_ = 0.00068, P_S_ = 1.28 × 10^-9^, P_P_ = 0.024). **i**, Bout lengths (number of stay trials before a switch) decreased for both high- and medium-reward patches (one-sided Wilcoxon rank-sum test: P_J_ = 8.94 × 10^-7^ and 1.64 × 10^-14^, P_W_ = 7.67 × 10^-10^ and 2.72 × 10^-4^, P_S_ = 3.05 × 10^-10^ and 3.03 × 10^-18^, P_P_ = 1.82 × 10^-5^ and 5.39 × 10^-8^). **j-l**, Behavioral metrics corresponding to g-i reproduced by simulations of the fitted behavioral model (same statistical tests as in g-i: P < 10^-5^ for all early-vs. late-learning comparisons). Error bars: 95% CI. **m**, Logistic regression coefficients predicting single-trial switch/stay decisions from model-estimated patch values (see Methods). Values of the current and alternative patches both significantly influenced rat choices, with opposite signs (P < 0.05 for coefficients of Value_curr. patch_ and maximal Value_alt. patch_ in each rat). We thus defined *switch value* as max(Value_alt. patch_) – Value_curr. patch_. **c, f-l**, Throughout the study, early learning comprised run sessions on day 1 of phase 2; late learning comprised run sessions on days 5-7, after the emergence of the new strategy, as indicated by a pronounced reduction in the estimated depletion factor.

In this task, a trial was defined as the interval between successive departures from two reward ports. Each port was assigned a nominal reward probability, p(R), of 0.2, 0.5, or 0.8, with one patch offering the highest average p(R) under a given assignment. On each visit to a port, reward delivery was determined by its p(R), and consecutive visits to the same port were never rewarded. During an initial training phase (1-2 weeks), the p(R) of all ports remained stable for blocks of 60 trials and changed covertly at block transitions within each session. As shown in our previous work^28^, rats learned to identify and exploit the patch with the highest reward probability and to adjust patch choices after the nominal p(R) changed.

We then introduced a second phase of the task that challenged animals to change how they learned from outcomes, that is, to go beyond purely outcome-based learning by incorporating an abstract reward rule. In this phase, the nominal p(R) was held fixed for each session while a new depletion-repletion rule altered the temporal structure of reward availability: if rats stayed within the same patch (i.e., alternated between its two ports), the p(R) of the visited port was discounted by 80% on the next visit, and restored only when they switched to a different patch (Fig. 1b). Each rat typically completed five 300-trial sessions per day, yielding ∼9500 total trials per animal. Nominal p(R) assignments were shuffled across sessions, allowing us to study both within-session reward learning and cross-session meta-learning of the new rule. Accordingly, all subsequent analyses focused on the second phase.

Inspection of behavioral patterns revealed that over days of training, rats adapted their foraging strategies as they gained experience with the depletion-repletion rule. In early-learning sessions, rats explored the three patches for extended periods and typically stayed within a given patch until reward was depleted, consistent with an action-selection strategy based on recent outcomes (Fig. 1c, top). In late-learning sessions, after an initial exploration phase, rats rapidly homed in on the high-reward patch (Fig. 1c, bottom; Extended Data Fig. 1a), often identifying the medium-reward patch as well. They also adopted a novel strategy: following exploitation of the high-reward patch, rats made brief visits to a lower-reward patch before returning to the high-reward one (Fig. 1c, bottom; Extended Data Fig. 1a). Remarkably, they commonly switched back to the high-reward patch even when their visits to the lower-reward patch were rewarded. These behavioral shifts were accompanied by increased overall reward rates, a hallmark of more adaptive reward learning (Extended Data Fig. 1b). This acquired strategy indicates the emergence of meta-learning—the moment-to-moment action policy was shaped by an internal model that, rather than relying solely on recent returns at each patch, forecasted impending depletion in the current patch and renewed availability elsewhere.

To quantify these reward-learning and decision-making patterns, we fit a statistical behavioral model to rats’ trial-by-trial choices and reward outcomes (Fig. 1d; see Methods). The model generated single-trial estimates of subjective port and patch values, enabling analysis of value-related computations associated with individual decisions (Fig. 1e). Specifically, it embedded a Bayesian form of recency-weighted reward learning, which is analogous to Q-learning but infers reward probability based on a Beta-Bernoulli distribution^33^. It was further augmented with a parameterized depletion rule that allowed obtained outcomes to deviate from the underlying reward probability. By fitting the model, we inferred a ‘depletion factor’ for each day that captured how strongly the depletion rule was incorporated into choice evaluation (Fig. 1f) and substantially improved prediction of rat choices in late days (Extended Data Fig. 1c). This factor was close to one during early learning, which corresponds to standard recency-weighted learning (i.e., no depletion). It declined to levels near the true discounting rate of 0.8 during late learning (Fig. 1f), suggesting that choices became increasingly aligned with the rule. Concurrently, the model estimated reduced ‘memory decay’ across learning (Extended Data Fig. 1d), consistent with more persistent memory of the underlying reward probability despite temporary fluctuations in reward experience.

We further validated the presence of meta-learning in each rat’s behavior by measuring changes in their decision tendencies and comparing these changes to simulations from the fitted behavioral model (Fig. 1g-l). Across learning, rats became more likely to choose the high-p(R) patch when switching from other patches and more likely to choose the medium-p(R) patch when leaving the high-p(R) patch (Fig. 1g, j). These shifts reflected knowledge of fixed nominal reward probabilities within each session. In addition, rats left a patch immediately after reward more frequently (Fig. 1h, k) and exhibited shorter bout lengths in both high-p(R) and medium-p(R) patches (Fig. 1i, l), congruent with internalization of the depletion-repletion rule.

The model’s ability to reproduce rats’ behavioral patterns motivated us to use its trial-by-trial value estimates to identify a decision variable predictive of rats’ choices. Choices were significantly influenced by both the value of the current patch and the maximal value of alternative patches (Fig. 1m, top). Their opposing contributions suggest that decisions were guided by a comparison of local and distant reward opportunities^34^, a decision variable we termed the ‘switch value’ (Fig. 1m, bottom; see Methods). Given its tight link to foraging behavior, we reasoned that the neural basis of this switch value (hereafter, ‘value’ for short) offers a mechanistic entry point for understanding how the brain shifts from a strategy of primarily caching recent reward history to one shaped by meta-learning of higher-order reward rules.

### Multiplicative coding of task structure and value at multiple levels of task abstraction

We then set out to understand how single mPFC neurons (n = 70-500 recorded simultaneously per day) encoded switch value and other task-relevant variables. As rats traveled between reward ports (defined as ‘journeys’), single-neuron activity showed three notable features: (i) temporal profiles that generalized across journeys sharing common task structures—such as goal progress and action sequence—irrespective of physical location (Fig. 2a); (ii) value modulation that appeared as a multiplicative gain on task-structure coding, with different neurons encoding value at distinct journey stages and at varying levels of abstraction (Fig. 2b); and (iii) value modulation across the mPFC population that collectively spanned the entire progression of each journey (Fig. 2b). We systematically characterized these encoding properties by fitting generalized linear models (GLMs) to single-neuron activity^35^ (see Methods). We compared regression coefficients of GLMs fitted separately for each journey to assess the generality of task-structure and value encoding across task conditions.

**Fig. 2.**
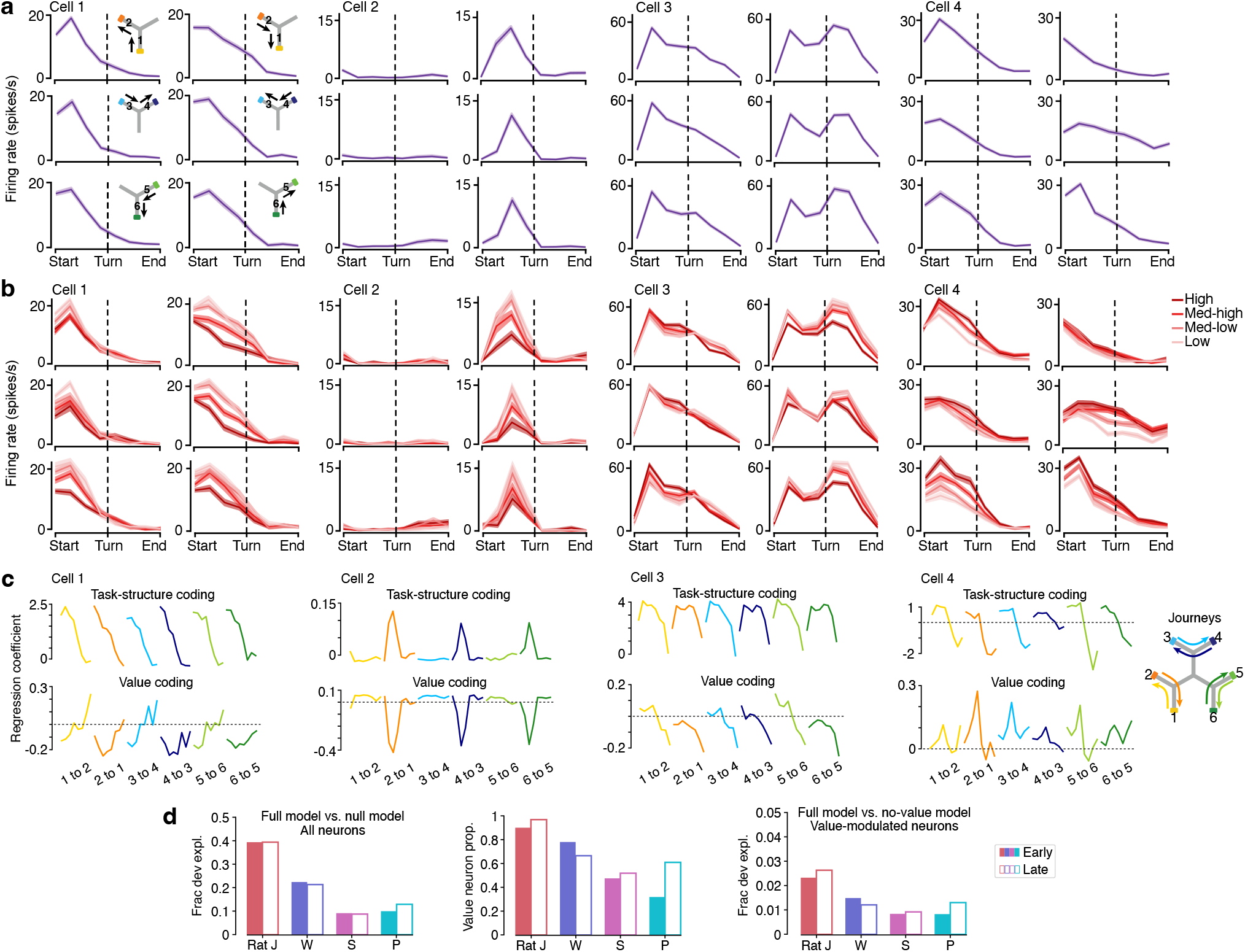
Mixed coding of task structure and value across task conditions in mPFC neurons. **a**, mPFC neurons were modulated by task structure such as goal progress, action, and patch identity. Shown are firing rates of four example neurons over 8 progression bins across 6 patch-stay journeys. Cell 1 exhibited consistent temporal profiles across all journeys. Cell 2 was selectively active during right-turn journeys. Cell 3 was active during both left- and right-turn journeys but showed action-dependent temporal profiles. Cell 4 showed broadly consistent temporal profiles across all journeys, with firing-rate ranges exhibiting specificity for patch 2 and patch 3. Inset: schematics of journeys associated with each firing profile. **b**, Value modulation appeared as a multiplicative gain on top of task-structure coding. For each neuron, trials within a journey were grouped into four quantiles based on their estimated switch value. Value-coding strength and sign depended on goal progress (highest degree of task abstraction), action, patch identity, and/or journey identity (lowest degree of abstraction). **c**, GLM regression coefficients for 8 progression bins per journey (top) and for 8 switch value × progression terms per journey (bottom). Same example neurons as in a, b. **d**, Summary of GLM results. Left: fraction of deviance explained by the full GLM compared to the null model in early- and late-learning sessions, averaged across all neurons per rat. Middle: proportion of value-modulated neurons in early- and late-learning sessions. Value-modulated neurons were defined as those for which the full model explained a significantly greater fraction of deviance than a reduced model lacking value × progression × journey terms (P < 0.01). Right: fraction of deviance explained by the value × progression × journey terms in early- and late-learning sessions, averaged across value-modulated neurons per rat. The same measurements across all days per rat are shown in Extended Data Fig. 2a to assess statistical significance.

We found that a substantial fraction of recorded mPFC neurons (30-90% per rat) showed significant modulation by value, with heterogeneous patterns of generality in both task-structure and value coding (Fig. 2c, d; Extended Data Fig. 2a): some neurons exhibited consistent regression-coefficient profiles across all stay journeys; others showed restricted generalization limited to journeys involving the same action (left vs. right turns) or within specific patches.

To determine the predominant level of value-coding generality for each neuron, we fit additional GLMs across journeys pooled by shared task structures and compared these model fits (see Methods). This approach allowed us to quantitatively classify neurons spanning a hierarchy of generalization. Consistent with the heterogeneity of cross-journey correlations of GLM value-coding coefficients, some neurons were best described by journey-specific models, some by action- or patch-dependent models, and others by progression-based models (Extended Data Fig. 2b).

Together, these results demonstrate that mPFC neurons possess diverse and hierarchically organized coding of task structure and value, ranging from broad generalization across journeys to condition-specific patterns. Importantly, the average encoding strengths across individual neurons remained stable between early and late sessions, showing no systematic changes over meta-learning (Fig. 2d, Extended Data Fig. 2c). This motivated us to investigate whether meta-learning instead altered how value was represented in the population through reorganization of neural dynamics.

### Generalized dynamical motifs for tracking, predicting, and updating values

Having observed that individual mPFC neurons encoded value in conjunction with task-structure variables including goal progress, action, and patch identity, we next asked how such heterogeneous mixed codes were organized at the population level. Because meta-learning in this task required internalizing the depletion-repletion rule and applying it across task conditions, we examined whether population activity expressed dynamical motifs (i.e., recurring patterns) that could serve as substrates for tracking, predicting, and updating switch value in different task contexts. Our goal here is to build intuition for the underlying structure of population dynamics, which we then quantify in subsequent analyses.

Given the generality of single-neuron coding, we first sought dynamical motifs that were shared across journeys. To avoid arbitrarily discretizing value estimates and presupposing value-coding structure, we instead grouped trials by their proximity to a patch switch. We then applied principal component analysis (PCA) to group-averaged neural population activity during the run period. This defined a ‘navigation subspace’ wherein the leading 6-7 PCs collectively explained >80% of the total variance (Extended Data Fig. 3). In this subspace, the most prominent structure was progress-tracking dynamics: neural trajectories traced out journey-invariant loops in the low-dimensional projection (Fig. 3a), reflecting the prevalence of goal-aligned sequential firing in single mPFC neurons (Fig. 2 and refs ^13,18,22^). Notably, along dimensions roughly orthogonal to these loops, trajectories were systematically displaced in a value-related order over successive trials leading up to a switch (Fig. 3a).

**Fig. 3.**
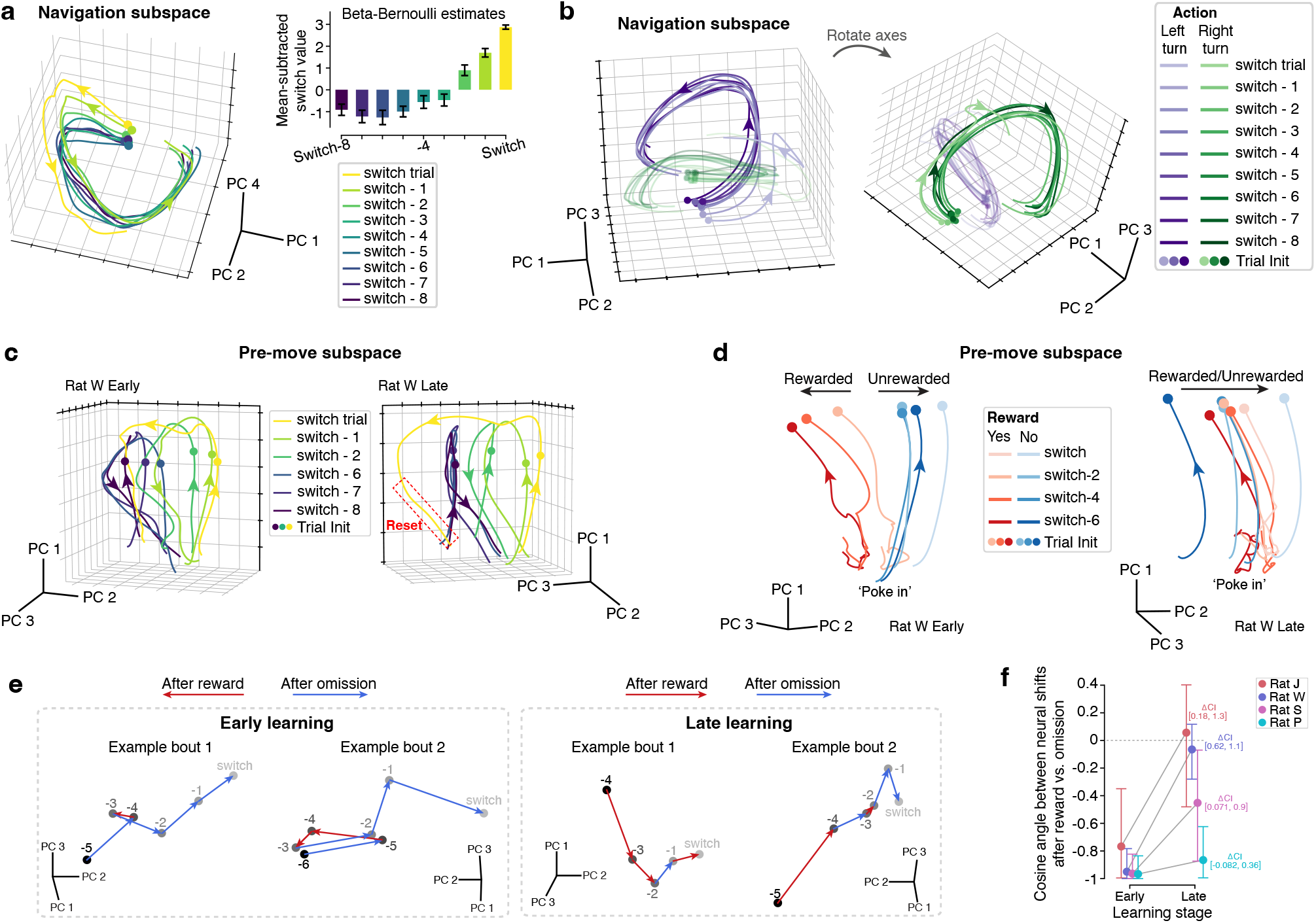
Dynamical motifs integrating task structure and value are reorganized with meta-learning. **a**, Run-time neural trajectories (0 to 3 s from trial initiation) projected into the leading PC subspace constructed from run-time neural activity (example day). Goal-progress dynamics dominated PCs 1 and 2. Approximately along an orthogonal axis (PC 4 in this example), the arranging order of these trajectories reflected increasing switch value. Inset: switch values computed from the Beta-Bernoulli estimates (mean-subtracted for clearer visualization of the gradient; error bars, standard deviation). *Before mean subtraction, switch values were consistently negative due to a patch-stay bias added to the current-patch value to capture the cost of switching* (see Methods). **b**, Run-time neural trajectories from left-turn (purple) and right-turn (green) trials leading up to a switch in the navigation subspace (example day). Within each action condition, trajectory displacement was systematically aligned with switch value, revealing action-specific variants of a common dynamical motif. **c**, Neural trajectories in the leading PC subspace constructed from pre-move neural activity (-0.8 to +0.2 s from trial initiation) formed a spiral motif (example rat). To cover both the end of value updating on the previous trial and the run period of the current trial, we projected neural data during the -0.8 to +3 s window (stay trials) and the -0.8 to +4.5 s window (switch trials) from trial initiation. After rats entered a new patch, the switch-trial neural trajectory (yellow) reset farther away from the stay trajectories in late learning (right) than in early learning (left). Red box highlights the reset trajectory during the final segment of a switch path. **d**, Neural trajectories in the pre-move subspace following reward vs. omission (example rat). To cover the value-updating period of the previous trial before initiation of the current trial, we projected neural data during the median outcome period (-5 to 0 s from post-reward trial initiation and -0.8 to 0 s from post-omission trial initiation). Note that among early-session trajectories (left), there was no switch trial right after reward. ‘Poke in’ denotes arrival at the port. **a-d**, Circles: trial initiation; arrowheads: neural trajectory direction. The PC axes and viewpoints for visualization were chosen to highlight value-related gradients. **e**, LFADS-denoised single-trial neural states at trial initiation after reward or omission in the pre-move subspace (example bouts). In early learning, neural states shifted in opposite directions after reward vs. omission. In late learning, these shifts aligned toward the switch direction (approximately the right side of PC 2 in the examples) after both outcomes. Numbers and grayscale of circles denote proximity to a switch. Red arrows: neural shifts after reward; blue arrows: shifts after omission. **f**, Cosine angles between neural state shifts following reward and omission (LFADS-denoised single-trial data, same as in e). Cosine angles in early learning were close to -1, indicating antiparallel shifts. They increased significantly in late learning for three rats (J, W, and S) and showed a weak upward trend in rat P, consistent with more aligned neural shifts. Shifts were computed along the two leading PC axes where spiral neural trajectories showed salient value-related gradients. Because these axes captured dominant variance in population activity rather than specifically targeting value-coding dimensions, other task variables may also contribute to the measured neural shifts. Error bars: 95% CI.

Similar value-displaced loops were found at finer levels of task generality—across journeys sharing actions (left vs. right turns), across journeys in the same patch, and even within individual journeys (Fig. 3b, Extended Data Fig. 4). Accordingly, mPFC population activity expressed a family of common dynamical motifs that jointly represented task structure and value. These motifs are well suited to provide a unified scaffold for integrating condition-specific task variables with global value estimates to flexibly inform foraging decisions in various contexts, thereby instantiating a generalized computational principle.

We next examined whether these structured population dynamics were sculpted by meta-learning in support of the rule-based strategy. Because neural variance in the navigation subspace was dominated by goal progress, we sought to extract additional dimensions that captured variance more strongly related to value. We therefore applied PCA to population activity during a pre-move time window (-0.8 to +0.2 s from trial start), a subset of the pause period when rats were stationary at reward ports and about to initiate the next movement (Extended Data Fig. 5). This yielded a ‘pre-move subspace’ into which we projected neural activity spanning both pause and run periods.

Within the pre-move subspace, we identified a spiral dynamical motif characterized by a salient value-related gradient across pre-switch and switch trials (Fig. 3c, Extended Data Fig. 6). Across pre-switch trials in both early- and late-learning sessions, neural trajectories progressed along the spiral in the switch direction, paralleling the ramp-up of switch value linked to patch depletion. On switch trials, these trajectories advanced further along the spiral before trial initiation.

A critical feature of this motif that changed markedly with meta-learning was how neural trajectories evolved as rats traversed to a new patch before any reward outcomes were revealed. During late learning when rats expressed the meta-learned strategy, switch-trial trajectories ‘reset’ prominently away from the switch direction, typically extending well beyond stay-trial trajectories or reaching levels comparable to early-stay states in the new patch (Fig. 3c, right; Extended Data Fig. 6, bottom). This anticipatory neural reset indicates a strong re-initialization of neural states prior to outcome delivery, consistent with inference about patch repletion. By contrast, the reset was moderate or absent in early learning (Fig. 3c, left; Extended Data Fig. 6, top): switch-trial trajectories either remained within the distribution of stay-trial trajectories rather than resetting further away, or showed minimal separation from switch states. The emergence of such augmented resetting suggests that task-state inference develops as a meta-learned computation.

In addition to pre-outcome state inference, we asked how the spiral motif mediated value updating during the outcome period and whether the updating mechanism, a crucial differentiator between recency-weighted and rule-based strategies^36^, changed with meta-learning. In early learning, neural trajectories in value-related dimensions exhibited clear outcome-dependent divergence after outcome delivery. Specifically, they unfolded in opposite directions following reward versus omission (Fig. 3d, left). Remarkably, in late learning, value-related neural trajectories progressed toward the switch direction across pre-switch trials regardless of preceding reward outcomes (Fig. 3d, right). This pattern is consistent with knowledge of depletion overriding purely outcome-driven value updating. To examine whether the altered updating process was also evident at the level of individual trials with distinct reward histories, we applied Latent Factor Analysis via Dynamical Systems (LFADS) to recover denoised single-trial dynamics (see Methods). Trial-by-trial shifts of neural states at trial initiation were nearly antiparallel after reward versus omission during early learning, but became more aligned toward the switch direction during late learning (Fig. 3e, f).

### Value-coding axes capture representational features of the dynamical motif

The PCA and LFADS trajectories pointed to systematic changes with meta-learning in the population dynamics supporting value computation. To directly quantify value-coding structure, we constructed cross-validated decoders of switch value from mPFC population activity. Given that PCA projections suggested generalized value representations across task conditions that were consistently intertwined with progression, we first targeted the most general case: the conjunctive coding of goal progress and value embedded in the spiral motif spanning both run and outcome periods. To this end, we pooled all stay trials and discretized each trial into eleven progression bins, including eight during run time and three during the outcome period. We then trained a separate value decoder for each bin using LASSO regression (see Methods).

These decoders explained a substantial fraction of the switch value variance (15-60%) throughout progression, with peak performance in pre-choice bins where value was most relevant to upcoming decisions (Fig. 4a, Extended Data Fig. 7a). Further, during the outcome period, decoding performance was higher for the updated value of the next trial than for the value of the current trial (Fig. 4a, Extended Data Fig. 7a), pointing to an active value-updating mechanism that bridged value representations in successive decision episodes (Extended Data Fig. 7b).

**Fig. 4.**
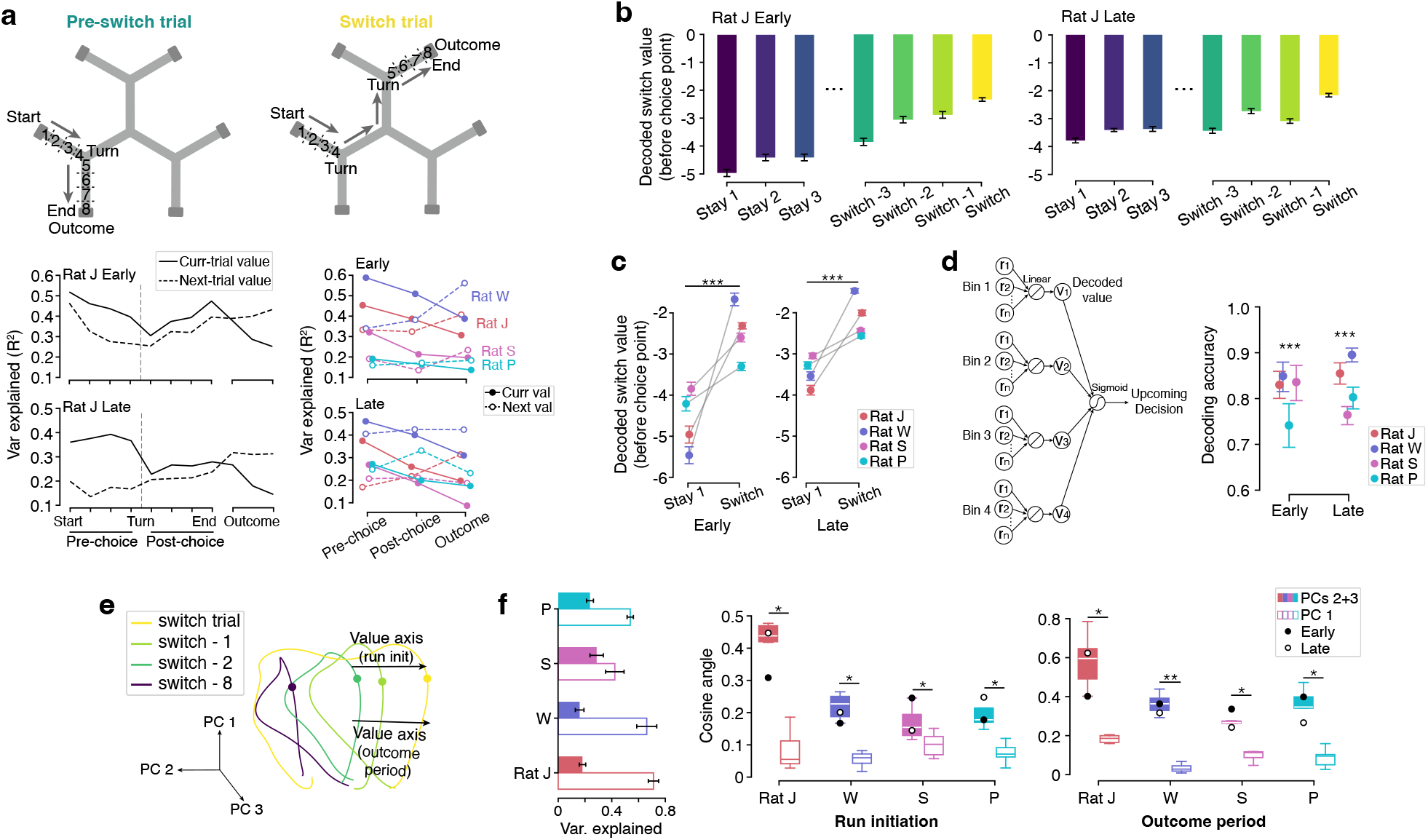
A linear value decoder identifies value-coding and value-updating dimensions. **a**, Decoder design and performance. Top: Schematic showing the progression-bin division of each trial used in value decoding. Switch trials were always held out as test data. Bottom left: Variance of switch value (estimated by the behavioral model) explained by the neural decoder in each progression bin (example rat, early-vs. late-learning sessions). Bottom right: Decoder performance averaged across pre-choice (bins 1-4), post-choice (bins 5-8), and outcome (bins 9-11) bins (early vs. late learning in all rats). During the pre-choice period, the decoder explained more variance of the current-trial value (filled circles) than of the next-trial value (hollow circles); during the outcome period, this pattern reversed. When decoding the current-trial value, performance was higher in pre-choice bins than in other bins. Statistical significance of these trends across all days and animals is tested in Extended Data Fig. 7a. **b**, Decoded switch value during the pre-choice period across early stay trials (Stay 1-3), late stay trials preceding a switch (Switch-3 to Switch-1), and the switch trial, shown for early- and late-learning sessions of an example rat. **c**, Decoded switch value increased significantly from the first trial (‘Stay 1’) to the final trial (‘Switch’) initiated within a given patch in both early and late learning (linear mixed effects models testing random slopes per animal: P_J_ = 1.23 × 10^-72^, P_W_ = 2.73 × 10^-75^, P_S_ = 1.78 × 10^-27^, P_P_ = 9.09 × 10^-17^ in early learning; P_J_ = 3.06 × 10^-105^, P_W_ = 2.29 × 10^-181^, P_S_ = 1.06 × 10^-52^, P_P_ = 8.42 × 10^-37^ in late learning), consistent with the displacement of neural trajectories in the PC subspace (Fig. 3a, c). **b, c**, Error bars: SEM across trials. **d**, Predicting decisions from neural value representations. Left: Schematic showing the construction of value and decision decoders. r_1_-r_n_: neural population activity. V_1_-V_4_: decoded switch value in progression bins 1-4. Right: Decoding accuracy of switch/stay decisions in early and late learning (bootstrap test for accuracy > 0.5: P = 0 in all rats). To avoid accuracy inflation due to imbalanced class counts, we measured the decoding performance for switch and stay decisions separately (Extended Data Fig. 7c) and took their mean. Error bars: 95% CI from bootstrapping. **e, f**, Alignment between value-decoding axes and PC dimensions. **e**, Value-decoding axes projected into the pre-move PC subspace on a representative day. **f**, Left: Across all days per rat, PC 1 explained more variance of the pre-move neural population activity than PCs 2+3. Right: Cosine similarity between PCs 1-3 of the pre-move subspace and the value-decoding axis of the first progression bin (‘run init’) or the outcome-period bin. Because spiral trajectories showed pronounced value-related displacement along PC 2, PC 3, or their combination (varying across days and animals), we computed the cosine angle between the value-decoding axis and the PCs 2+3 plane (filled boxes). Boxes include all days of each animal to assess statistical significance; early (filled circles) and late (hollow circles) days are highlighted to demonstrate stability of this alignment. By contrast, alignment with PC 1 was significantly weaker (empty boxes), supporting the idea that the correlation between the value-decoding axis and PCs 2+3 was not due to chance (one-sided paired Wilcoxon signed-rank test: P_J_ = 0.016, P_W_ = 0.016, P_S_ = 0.031, P_P_ = 0.031 during run initiation; P_J_ = 0.016, P_W_ = 0.0078, P_S_ = 0.016, P_P_ = 0.031 during the outcome period).

Having established robust decoding performance, we next examined how neural value representations evolved across trials and related to decision-making. Because value readouts were largely stable across pre-choice bins (Fig. 4a, Extended Data Fig. 7a), we averaged them to obtain a single pre-choice value per trial. We observed that, in both early and late learning, decoded switch value ramped up across pre-switch trials (Fig. 4b), yielding significantly higher values on switch than on stay trials (Fig. 4c). This graded increase mirrored the arrangement of neural trajectories in the spiral motif. Moreover, decoded switch value in pre-choice bins accurately predicted single-trial switch/stay decisions in all rats, highlighting the behavioral relevance of the neural value representations (Fig. 4d, Extended Data Fig. 7c). Notably, the value-decoding axes during both run-initiation and outcome-period bins aligned with directions in the PC subspace along which neural trajectories were displaced according to value (Fig. 4e, f). Together, the neural decoders provide an explicit readout of value computations embedded in the spiral dynamical motif.

Finally, we evaluated whether such value computations were expressed consistently across multiple levels of generality, e.g., for journeys within the same patch, for left-versus right-turn journeys, and for each individual journey. We fit decoders at each generality level, all of which achieved reliable performance (Extended Data Fig. 7d; see Methods). This decoding robustness aligns with the coexistence of value-related dynamical motifs across task conditions and further corroborates the generalizability of the underlying value computations.

### Changes in the value-coding dynamics over meta-learning

The neural value decoder provided a direct means to quantify the reorganization of mPFC population dynamics associated with meta-learning, as indicated by the reshaped spiral trajectories in late sessions. We therefore examined longitudinal changes in neural value representations as animals internalized the depletion-repletion rule. Motivated by the pronounced resetting of spiral trajectories in late learning (Fig. 3c), we first asked whether neural value representations exhibited similar resets. To this end, we compared decoded switch value at patch departure (‘switch-out value’) with that at the next patch re-entry before outcome delivery (‘switch-in value’) for the same patch. A reduction from switch-out to switch-in value reflects neural value resetting and provides a measure of ‘neural value repletion’. Guided by the behavioral model metrics, we expected moderate neural resetting early in training due to memory decay of cached values toward baseline (Extended Data Fig. 1d, Fig. 5a), and stronger resetting later due to incorporation of the repletion rule (Figs. 1f, 5a).

**Fig. 5.**
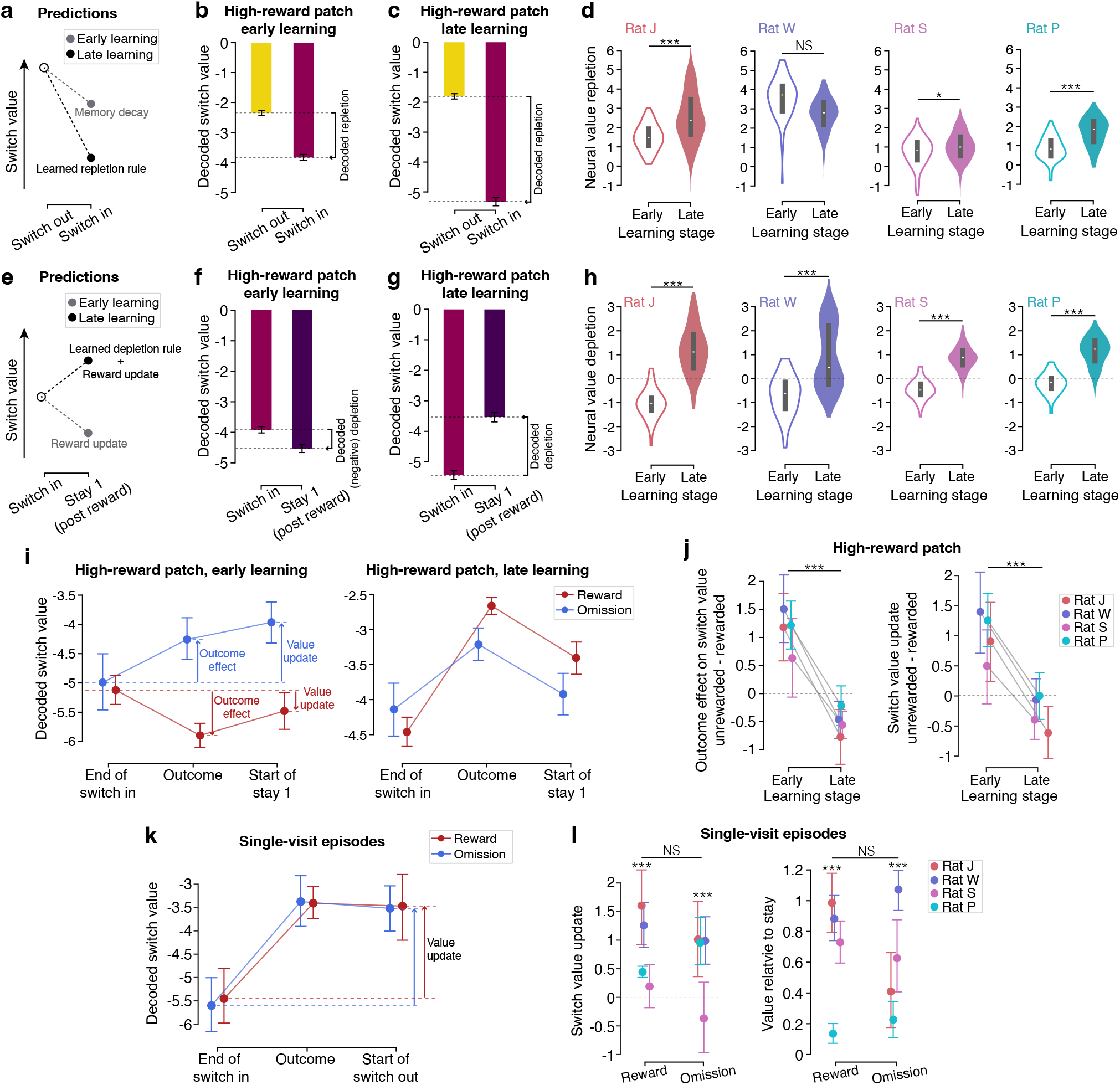
Changes in neural value representations reflect the meta-learned strategy. **a**, Conceptual predictions based on neural state-space trajectories (Fig. 3c) and behavioral model estimates (Fig. 1f, Extended Data Fig. 1d). Early learning was expected to show moderate resetting of decoded switch value due to decay toward baseline, whereas late learning was expected to show substantial resetting driven by incorporation of the repletion rule. **b, c**, Decoded switch value during the pre-choice period (mean of bins 1-4; see schematic in Fig. 4a) of the switch-out trial and the post-choice period (mean of bins 5-8) of the switch-in trial in the high-reward patch (example rat). Error bars: SEM. **d**, Neural value repletion, quantified as *switch-out value – switch-in value*, increased significantly on late-learning days for three rats (J, S, P); rat W showed strong neural repletion in both early and late learning, likely reflecting pronounced memory decay and/or rapid rule learning on day 1 (one-sided Wilcoxon rank-sum test: P_J_ = 5.65 × 10^-11^, P_W_ = 0.99, P_s_= 0.029, P_P_ = 2.45 × 10^-12^; linear mixed effects model testing fixed slope: P = 0.00034). **e**, Conceptual prediction that decoded switch value decreases after a rewarded switch-in trial in early learning due to recency-weighted value updating, but increases in late learning once the depletion rule overrides reward update. **f, g**, Decoded switch value during the post-choice period of the switch-in trial and its subsequent stay trial in the high-reward patch (example rat). Error bars: SEM. **f**, Early learning: decoded switch value decreased after a rewarded switch-in trial, which defined negative neural value depletion. **g**, Late learning: decoded switch value increased after a rewarded switch-in trial, reflecting substantial neural value depletion. **h**, Neural value depletion, quantified as *first-stay switch value – switch-in value*, increased significantly in late learning for all rats (one-sided Wilcoxon rank-sum test: P_J_ = 1.40 × 10^-19^, P_W_ = 1.51 × 10^-13^, P_s_= 1.31 × 10^-23^, P_P_ = 3.76 × 10^-17^). **i**, Reward and omission updated neural value representations in opposite directions in early learning (left) but not in late learning (right). One example rat. We defined (i) ‘outcome effect’ as decoded switch value during the outcome period minus that of switch-in bin 8, and (ii) ‘value update’ as decoded switch value of first-stay bin 1 minus that of switch-in bin 8. **j**, Left: In all rats, the difference between outcome effects following unrewarded vs. rewarded trials decreased significantly in late learning (linear mixed effects models testing random slopes per rat: P_J_ = 1.70 × 10^-5^, P_W_ = 3.62 × 10^-7^, P_S_ = 2.02 × 10^-4^, P_P_ = 1.17 × 10^-5^; fixed slope across all rats: P = 2.88 × 10^-19^). Right: A similar reduction was observed for the difference in value updates (linear mixed effects models testing random slopes per rat: P_J_ = 4.87 × 10^-4^, P_W_ = 7.84 × 10^-4^, P_S_ = 0.011, P_P_ = 9.12 × 10^-4^; fixed slope across all rats: P = 1.59 × 10^-9^). Error bars: 95% CI. **k**, Decoded switch value following single visits increased regardless of outcomes (example rat). Switch value update was defined as in i, except that the first-stay trial was replaced by the switch-out trial. **l**, Neural signatures of meta-learning during single-visit episodes in late learning. Left: Switch value updates were significantly greater than 0 after both rewarded and unrewarded single visits (one-sided Wilcoxon signed-rank test across rats: P_rewarded_ = 1.75 × 10^-13^, P_omission_ = 2.56 × 10^-6^), with no significant difference between them (linear mixed effects model testing fixed slope: P = 0.24). Right: Decoded switch value on switch-out trials following both rewarded and unrewarded single visits was significantly greater than that on stay trials in the same patch (one-sided Wilcoxon signed-rank test across rats: P_rewarded_ = 1.98 × 10^-23^, P_omission_ = 3.76 × 10^-18^), with no significant difference between reward outcomes (linear mixed effects model testing fixed slope: P = 0.72). A total of 147 rewarded and 115 unrewarded single-visit trials contributed to the analysis across all rats. Error bars: 95% CI.

In line with these predictions, we observed that greater neural value repletion emerged with meta-learning in three out of four rats (Fig. 5b-d). In early learning, switch-in value showed modest reductions relative to switch-out value (Fig. 5b, d). By contrast, in late learning, switch-in value dropped substantially below switch-out value (Fig. 5c, d). This enhanced neural repletion, occurring prior to outcome delivery, reflects anticipatory inference about replenishment of the newly entered patch rather than updating from direct reward experience. Importantly, switch trials were always held out of decoder training, ensuring that these neural effects cannot be attributed to structure imposed by the behavioral model.

Beyond pre-outcome inference, neural dynamics during the outcome period suggested that meta-learning also reshaped value updating in accordance with the depletion rule (Fig. 3d-f). This prompted us to hypothesize that reward should drive decoded switch value downward in early learning but exert a reduced or reversed effect once the depletion rule was internalized (Fig. 5e). To test this, we compared switch-in value with the decoded value on the subsequent stay trial within the same patch (‘first-stay switch value’), restricting analyses to rewarded switch-in trials (Fig. 5f-h). Consistent with recency-weighted value updating, decoded switch value decreased after reward early in training (Fig. 5f, h; Extended Data Fig. 8a). In late-learning sessions, however, first-stay switch value increased relative to switch-in value despite reward receipt (Fig. 5g, h; Extended Data Fig. 8b). This pattern implies that updating was primarily guided by inference about imminent depletion of the current patch.

To further dissect the effects of outcomes on neural updating, we examined how neural value representations evolved following rewarded and unrewarded switch trials. We quantified neural value update as the change in decoded switch value from the end of a switch trial to the start of the subsequent stay and then compared updates after opposite outcomes (Fig. 5i). We found that, during early learning, decoded switch value was updated more positively after omission than after reward, consistent with outcome-driven updating. By contrast, reward and omission no longer produced opposing updates during late learning (Fig. 5i, j; Extended Data Fig. 8c, d). This convergence of post-outcome updates signals a transition toward rule-based value updating.

The decoupling of value updating from immediate reward outcomes was most evident in an extreme case of the meta-learned strategy: in a subset of late-learning sessions, rats visited a lower-reward patch for a single trial and then returned to the replenished high-reward patch, even when that visit was rewarded (Fig. 1c, bottom; Extended Data Fig. 1a, bottom). This strategy exploited repletion and would generally not be expected to arise under baseline recency-weighted reward learning. These single-visit episodes thus presented a unique opportunity to distinguish purely reward-driven from rule-based neural updating mechanisms.

We found that changes in decoded switch value from switching in to switching out were similar for rewarded and unrewarded single visits (Fig. 5k). In both cases, the updated values at the start of the subsequent switch-out trials lay closer to the switch boundary (Fig. 5l, left; Extended Data Fig. 8e). Further, these switch-out values were consistently higher than values decoded for stay trials in the same patch (i.e., when rats remained in the patch after switching in), regardless of the single-visit reward outcomes (Fig. 5l, right; Extended Data Fig. 8e). These results, together with enhanced neural value repletion and depletion, demonstrate that during meta-learning (here, internalizing the depletion-repletion rule), frontal network dynamics undergo systematic reorganization associated with pre-outcome value inference and rule-guided value updating, thereby aligning decisions with the latent structure of the environment.

## Discussion

Learning requires updating internal representations based on experience, and meta-learning refines these updating processes to better match the statistical structure of the environment. Our work provides a direct window into how prefrontal cortical dynamics and computations are modified during meta-learning. We developed a spatial foraging task in which animals transitioned from learning stable reward contingencies to incorporating a higher-order depletion-repletion rule. We then investigated value representations in both single mPFC neurons and population activity. Value coding in individual neurons was mixed with task-structure coding typically through gain modulation, which, at the population level, was organized into dynamical motifs that tracked goal progress, option values, and other task features. These motifs generalized across task conditions, suggesting that mPFC population dynamics implement value computations in a condition-flexible manner. Notably, as animals meta-learned the depletion-repletion rule, value representations in the neural population changed markedly: neural dynamics came to express rule-guided inference of pre-outcome value and rule-based value updating, overriding purely outcome-driven computations. These findings indicate that meta-learning reorganizes mPFC dynamics underlying value tracking, updating, and prediction in support of more adaptive decision-making in complex, changing environments.

Meta-learning has been reported across a wide range of behavioral paradigms^3–6,11^, and converging evidence implicates PFC in acquiring abstract task rules and refining learning strategies over experience^9,10,37,38^. At the single-neuron level, prior studies have shown PFC coding properties that reflect the learning of abstract rules^39,40^. More recently, population-level analyses have revealed that as animals become proficient at reward learning in a two-arm bandit task, value representations in orbitofrontal cortex (OFC) converge across days toward a stable coding axis^10^. Building on these insights, our results present several primary advances:

First, we identified a spiral dynamical motif and associated single-neuron mixed selectivity that conjunctively represent task structure and value. This motif resembles value-coding cylindrical dynamics observed in retrosplenial cortex during model-free reinforcement learning^41^, suggesting that such motifs may constitute a broader class of population dynamics used across brain regions. It is worth noting that the retrosplenial motif has been attributed primarily to demixed, persistent coding of reward history, whereas the motif described here arises from mixed coding of value and task structure. Despite this distinction, the common expression of similar motifs points to a population mechanism for generalization, in which dynamical structures are reused to instantiate equivalent computations in different contexts. Critically, we found that meta-learning selectively reshaped population activity along value-coding dimensions while preserving accurate representations of other variables, including goal progress, action, and patch identity. This specific reorganization illustrates how meta-learning can flexibly adapt reward-learning algorithms without disrupting the broader representational scaffold, reconciling robustness with flexibility in PFC computation.

Second, because our task incorporated a depletion-repletion rule, animals learned to fundamentally revise their expectations about future reward availability rather than relying solely on recent outcomes. Correspondingly, during late learning, neural value representations exhibited a pronounced reset as animals re-entered a previously depleted patch, even before reward outcomes were revealed. This anticipatory reset indicates that mPFC dynamics came to encode rule-based inference about value repletion beyond merely caching reward history. While previous studies have established a critical role for mPFC in rule-based prediction^24,42^, our findings further demonstrate that such predictive representations can emerge through systematic, meta-learning-dependent changes in population activity.

Third, meta-learning also altered how reward outcomes influenced both behavioral policies and neural value updating. Behaviorally, animals shortened their stays within patches and increasingly left after a rewarded visit, marking a departure from a win-stay strategy. This shift was mirrored in mPFC population activity: early in learning, rewards consistently lowered neural representations of switch value, whereas later in learning the decoded switch value increased sharply even after reward receipt. Importantly, such reduced sensitivity to immediate outcomes did not reflect an absence of reward information. Instead, animals continued to use reward history to estimate repleted patch values and distinguish high-reward from lower-reward patches. Consequently, value updating was no longer well described by simple recency-weighted integration of wins and losses; rather, the directional impact of individual trials on value expectancy was governed primarily by the depletion rule. This selective modification of outcome influence underscores the precision with which meta-learning reshapes value-updating computations.

Our work bears directly on a key hypothesis from computational studies: that neural network dynamics and synaptic plasticity operate as complementary learning mechanisms across timescales, with the former implementing fast reward-learning algorithms and the latter mediating slower meta-learning processes that coach those algorithms^9–11^. This perspective posits that synaptic plasticity shapes the dynamical landscape across sessions, whereas moment-to-moment value updating unfolds through evolving activity states. Although synaptic mechanisms were not directly probed, our results provide empirical support for this framework by demonstrating that meta-learning sculpts mPFC population dynamics underlying value- and decision-relevant computations during the emergence of a new learning strategy. In particular, the dynamical motif we identified integrates instantaneous reward learning with internalized task knowledge, establishing a direct link between evolving neural computations and behavioral strategy transitions. Such an architecture is well suited to adapt reward learning when value updating departs from classic recency-weighted algorithms, as in the present task. Reorganization of neural dynamics may therefore endow decision-making with greater flexibility and foresight—animals can adjust expectations about future states and select actions based on an internal model of task rules rather than integrating recent outcomes under the assumption of stable reward probabilities.

Finally, our results place mPFC dynamics within a broader framework of meta-learning that engages distributed brain regions. Other PFC subregions, including OFC^10,25,43,44^, as well as subcortical regions such as the hippocampus and basal ganglia, have also been implicated in reward learning and meta-learning^45–50^, yet how their contributions are differentiated and coordinated remains an open question^9,48^. Our findings suggest that structured reorganization of population dynamics may represent a general mechanism through which these regions internalize task rules and modify learning algorithms without necessitating ongoing synaptic modifications.

## Methods

### Animals

All animal procedures were approved by the Institutional Animal Care and use Committee (IACUC) at University of California, San Francisco. Five adult male Long-Evans rats (Charles River Laboratories, 450-650g) were each pair-housed in a temperature- and humidity-controlled environment on a 12-hour light-dark cycle with lights on from 6 AM to 6 PM. Rats had *ad libitum* access to standard rat chow and water. Prior to behavioral training, rats were transitioned to single housing and food-restricted to maintain approximately 85% of their baseline weight. Note that one animal was excluded from analyses because of a different training regime and a poor fit of the behavioral model, making it difficult to make clear statements about value coding comparable to other animals.

### Behavioral pre-training

All behavioral tasks were controlled via custom code written in Python and Statescript in combination with an Environmental Control Unit (SpikeGadgets). The pre-training procedure was the same as described in our previous work^28^. Briefly, rats learned to run back and forth on a linear track for liquid food (sweetened evaporated milk) dispensed by reward ports at each end of the track. Upon entry of a port, 100 µL reward was delivered by a syringe pump immediately, gated by an infrared beam that rats learned to poke their noses into. The pre-training period was terminated when rats were able to gain at least 30 rewards in a 15-minute session. Pre-training took place in an environment fully separate from the spatial foraging task environment. Rats were subsequently returned to *ad libitum* food access prior to surgical implantation.

### Neural implant

Custom hybrid implants contained 24 independently movable tetrodes (California Fine Wire and Kanthal) and a 3D printed tetrode drive body (PolyJet HD, Stratasys) to target the hippocampus in each hemisphere. Details about the tetrode system can be found in our previous work^28^. Implants also contained a custom headstage (SpikeGadgets) coupled with a stacking set of up to four 128-channel polymer probes^29^ (Lawrence Livermore National Labs) to target the frontal cortex in each rat. The hybrid implant was protected in a 3D-printed enclosure. Drive and probes were sterilized before surgical implantation.

Both tetrodes and probes were implanted stereotaxically under sterile conditions. In anesthetized rats, cannulae were implanted above dorsal hippocampus and polymer probes were implanted in mPFC and OFC. mPFC probes were implanted with shanks (four shanks per probe) parallel to midline (+/-2.2 to 3.2 AP, +/-0.5 to 1.2 ML, and -3.2 to -4.3 DV relative to Bregma; 0-11 degrees lateral tilt in coronal plane). Only mPFC data have been analyzed in this study. A stainless-steel ground screw was implanted over the cerebellum to serve as a global reference. Titanium screws were placed in the skull to strengthen the implant and secured with Metabond (Parkell) and dental cement (Henry Schein).

After full recovery from surgery, rats were food deprived to 85% of their baseline weight and reintroduced to the linear track (described above), which refamiliarized them with obtaining reward on a track and prepared them for running with the implant. Two rats had additional prior experience on a fork maze^22^. This post-surgery re-training procedure ensured that rats were motivated enough to begin the spatial foraging task.

### Spatial foraging task with reward depletion and repletion

Before performing the main task in this study, all rats had completed 1-2 weeks of training on a no-depletion version of the spatial foraging task^28^ (phase 1). In both versions of the task, animals navigated in a maze containing three radially arranged Y-shaped ‘foraging patches’ (Fig. 1a). Each patch consisted of a central path that bifurcated into two subpaths and terminated in reward ports, resulting in two ports per patch and six ports in total. Paths and subpaths (‘track segments’) measured 6.5 cm in width and 53 cm in length. All central paths intersected at 120 degrees, such that the track had three-fold symmetry. The maze was made of black acrylic (TAP Plastics) with 3-cm-high walls that allowed rats to see distal spatial cues.

Each experimental day began with transferring the rat from its home cage to a rest box located in the same room as the main task maze. The first run session started after a 20-minute baseline rest session. In the first training phase (no-depletion task), rats performed seven to eight ∼20-min run sessions on each day (180 trials/session over three blocks). We then introduced the reward depletion-repletion rule. In the second phase, rats typically completed five ∼30-min run sessions per day (∼300 trials/session). In both phases, run sessions were interleaved with rest sessions of at least 30 minutes each. Rest sessions helped to maintain stable motivation by providing breaks throughout each day. For the depletion-repletion task, behavioral and neural data were collected over 1-2 weeks per rat. Throughout this study, early learning was defined as all run sessions on the first day of phase 2 prior to substantial rule acquisition, whereas late learning comprised all run sessions from days 5-7 after the new behavioral strategy had emerged, as indicated by a pronounced reduction in the depletion factor estimated by the behavioral model (see **Behavioral Beta-Bernoulli model** below).

During each run session, a rat was first placed at the track center facing a consistent direction, and then navigated freely in the environment under fully automated task control by Trodes software. Nominal reward probabilities p(R) of 0.2, 0.5, or 0.8 were assigned to each of the six reward ports (Fig. 1a). The two ports within a patch could share or differ in p(R), but the highest possible combination of p(R) was 0.5 and 0.8 (never 0.8 and 0.8). A trial was defined as the interval from exiting one reward port to exiting a different port (consecutive visits at the same port were never rewarded). Therefore, each trial consisted of (i) a run period between two distinct ports and (ii) an outcome period at the chosen port, during which the rat either received a 100 µL reward or experienced an omission. In phase 1 (no-depletion task), p(R) of each port remained stable within blocks of 60 trials and changed covertly at block transitions during a run session. During phase 2 (depletion-repletion task), each run session contained one fixed set of p(R) that was changed and counterbalanced across sessions. Additionally, p(R) was dynamically modulated by rats’ choices based on a depletion-repletion rule: if a rat alternated between the two ports within a patch on consecutive trials, p(R) of the visited port was discounted by 80% on the next visit and continued to deplete with repeated alternation; p(R) reset to its nominal value only when the rat switched to a different patch (Fig. 1b).

All sessions took place in a dimly lit room (2.4 m by 2.9 m) with black distal spatial cues on white walls. A ceiling-mounted overhead camera (Allied Vision) centered over the maze recorded rat behavior at 30 frames/second and was synchronized via Precision Time Protocol with all other signals recorded on Trodes software (SpikeGadgets).

### Histology

At the conclusion of behavioral experiments, rats were anesthetized with isoflurane, and small electrolytic lesions were made in the hippocampus. One day later, animals were anesthetized and transcardially perfused with 4% paraformaldehyde. Brains were fixed in situ overnight, after which brains were extracted from the skull. Brain fixation continued for two days. After cryoprotection in 30% sucrose for 5-7 days, brains were blocked and sectioned with a cryostat into 50-100 µm sections. mPFC sections were labeled with NeuroTrace 435/455 nuclear counterstain and glial fibrillary acidic protein GFAP (mouse anti-GFAP with A594 secondary antibody) to visualize probe implant sites (Extended Data Fig. 9).

### Data processing and analyses

All data processing and analyses were carried out in Python and Julia within the Spyglass pipeline to support reproducibility and data sharing^28,51^.

#### Spike sorting and curation

We performed spike sorting using MountainSort4, an automated spike sorting algorithm^22,29^. For 128 channel polymer probe recordings in PFC, all channels were referenced against a channel on the same probe with no spikes and bandpass filtered between 300 and 6000 Hz. Artifacts were identified and removed when voltage exceeded 1500 mV on over three quarters of channels of a probe shank. Spiking units were identified in neighborhoods of channels within 115 microns of each other. All spike sorting was applied to concatenated run and rest sessions within a day. After spike sorting, an automated round of curation was performed on the units identified by MountainSort4 using metrics calculated in SpikeInterface^52^. Units with metrics exceeding any of the following thresholds were rejected: a nearest neighbor noise overlap of 0.03, an inter-spike interval violation ratio of 0.0025, and a time shift in the waveform peak of two samples^22^. Then manual curation was performed on the remaining units to further improve the identification of single units and removal of artifacts. This process involved visual inspection to achieve two major objectives. First, different units whose cross-correlogram and waveform metrics satisfied criteria of belonging to a single neuron (yet with spikes incorrectly split during spike sorting) were merged. Second, individual units whose auto-correlograms corresponded to more than one neuron (i.e., multi-unit activity) were excluded. After spike sorting and curation, units with an average firing rate below 0.5 Hz within a given day were excluded. All remaining units were included in downstream analyses.

#### Behavioral Beta-Bernoulli model

To estimate the rats’ expected belief about the value of each of the six ports, we used a Beta-Bernoulli model, which estimates the likelihood of any possible value by tracking past successes (rewards) and failures (omissions) at that port. This infers not only the expected value of a given port (analogous to traditional Q-learning), but also the probability of the port having *any* possible underlying success rate. In fact, this model provides an optimal estimate of the underlying success rate given the number of past successes and failures, assuming that success rate is constant in time.

The Beta-Bernoulli model tracks port *i* with two variables, *α*_*i*_ and *β*_*i*_, which correspond to (with some adjustment, detailed below) the respective number of successes and failures at port *i*. Alternatively, the model can be re-parameterized via a mean and standard deviation *µ*_*i*_ and *σ*_*i*_, which eliminates the need to maintain a count of past events, and permits a simple neural implementation.

### Port Q value estimation

To estimate the Q value of any given port, the expected value of port *i* is given by 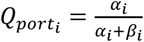, providing a simple estimate to compare ports.

### Patch Q value estimation

On each trial, the Q value of the current patch (if the rat were to stay) was computed as a weighted average of its two ports, *Q*_*patch*_ = *γ* * *Q*_*upcoming*_ + (1 − *γ*)*Q*_*past*_ where *Q*_*past*_ denotes the Q value of the port the rat departed from and *Q*_*upcoming*_ denotes the Q value of the port the rat was heading toward. For alternative patches, *Q*_*patch*_ was defined as the mean of the two port Q values within each patch.

### Learning rule

At the start of each session, *α*_*i*_ and *β*_*i*_ of each port were initialized to 1. After each port visit, the model was updated with the following rule:

After a reward at port *i*:

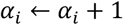

After an omission at port *i*:

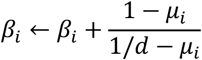

where *µ*_*i*_ is the expected value of port *i* given by 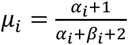, and *d* is the current estimation of reward depletion, beginning at 1 and multiplied by a *depletion factor* at each re-visit to the same port. Note that the above role departs slightly from a standard Beta-Bernoulli model where *β*_*i*_ would increment by 1 on an omission. This modification correctly accounts for the loss of information due to depletion. Briefly, consider the two extreme cases of *d*. Upon the first visit to a port, *d* = 1, and the above update rule reduces to the standard rule of *β*_*i*_ ← *β*_*i*_ + 1. At the other extreme, as *d* approaches 0 with repeated visits, the reward probability approaches 0, and the resulting update of *β*_*i*_ ← *β*_*i*_ presents the fact that a negative outcome provides minimal information about the underlying reward rate. The *depletion factor* was estimated for each day from rat choices and reward outcomes on each trial.

### Forgetting

To account for the possibility of forgetting when rats did not visit a port for a while, *α*_*i*_ and *β*_*i*_ were decayed towards 1 after each trial in which port *i* was not chosen:

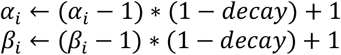

#### Behavioral choice model

To connect the value learning model to choices and assess our ability to capture animal behavior, we computed the likelihood of patch choice on every trial as the *softmax* of the three integrated patch values, given by:

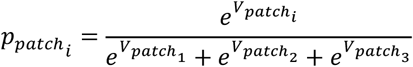

where the integrated value at the current patch was given by:

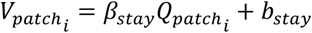

The value of the patch clockwise from the current patch was given by:

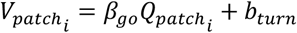

The value of the remaining patch was given by:

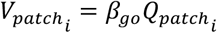

where 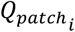 is the patch Q value estimated by the Beta-Bernoulli model as described above.

The inverse temperature parameters, *β*_*go*_ and *β*_*stay*_, determined how sensitive choice behavior was to differences in expected reward: high temperatures indicate high sensitivity to reward, while low temperatures indicate insensitivity. To allow for differential sensitivity to reward estimates at the current patch versus alternative patches, we fit two independent softmax temperatures, *β*_*stay*_ for the current patch, and *β*_*go*_ for the two alternative patches. The term *β*_*go*_ reflected a fixed cost to changing patches due to the time and distance to reach a new patch (as opposed to the reward sensitivity fit by *β*_*stay*_ that reflected a tendency to stay in the current patch).

To incorporate tendencies of the animals to prefer to turn in certain directions when changing patches, we included a *turn bias b*_*turn*_, an offset consistently applied to the patch to the left of the animal’s current position. If the rat was in patch A, this was added to patch B. If the rat was in patch B, this was added to patch C, and if the rat was in patch C, this was added to patch A.

On trials where the animal switched patches, likelihood of port choice was modeled as a subsequent softmax between the two ports in the new patch, with a separate softmax temperature *β*_*port*_ applied to the per-port Q value estimates:

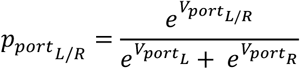

The integrated value of the left port was given by:

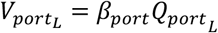

And the value of the right port was given by:

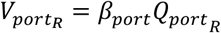

#### Behavioral model fitting procedure

For the Beta-Bernoulli model, we optimized the free parameters (*including γ, depletion factor, decay, β*_*go*_, *β*_*stay*_, *β*_*port*_, *b*_*stay*_ *and b*_*turn*_) of the model by embedding each of them within a hierarchical model to allow parameters to vary from day-to-day. These day-level parameters were themselves modeled as arising from a population-level Gaussian distribution over days, separately for each rat. We estimated the model for each rat to obtain optimal day- and rat-level parameters that minimized the negative log likelihood of predicting the actual choice using an expectation maximization (EM) algorithm with a Laplace approximation to the day-level marginal likelihoods in the M step^53^.

#### Definition of switch value

We defined the *switch value* on each trial as the difference between the highest-valued alternative patch and the current patch:

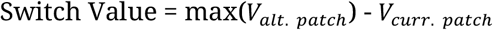

where patch values 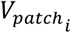 were estimated as described above. Larger switch values therefore indicate greater relative incentive to leave the current patch. Note that switch values were consistently negative because the behavioral model added a patch-stay bias to the value of the current patch to capture the cost of patch switching.

#### Generalized Linear Models (GLMs) for single-unit coding of task structure and value

We quantified how individual mPFC neurons encoded task structure and switch value using a previously established Poisson GLM framework^35,54^. In this framework, neuronal spikes are modeled as a Poisson process whose time-varying rate is a log-linear function of covariates (task and behavioral variables) during each regression bin. Because trials were self-initiated and varied in duration, we discretized each of the six stay journeys into eight goal-progress bins rather than using fixed-duration temporal bins (i.e., 48 journey × progression bins total). For each neuron, we computed spike counts within each progression bin on each trial. We then defined sets of covariates to capture (1) goal progress in each journey using a binary indicator that represents the presence or absence of the rat in a specific goal-progress bin, and (2) the interaction between goal progress and switch value:

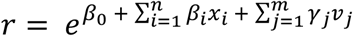

where *r* denotes a vector of firing rates for one neuron over *T* time points, *x*_*i*_ is a binary vector indicating whether an animal was in progression bin *i* (out of *n* bins total) at time *t*, and *ν*_*j*_ is a vector indicating the behaviorally-inferred switch value when the animal was in progression bin *j* (out of *m* bins total). To assess how broadly each neuron’s value coding generalized across task conditions (Extended Data Fig. 2b), we used the same basic GLM structure and retained the journey × progression bins (*x*_*i*_ for *i* = 1, 2,… 48), but adjusted the set of progression-value interaction terms (*ν*_*j*_) to reflect different condition groupings:

- Patch-level GLM: 24 patch ID × progression value terms
- Action-level GLM: 16 action × progression value terms
- Progression-level GLM: 8 goal-progress value terms

To assess the strength of task-structure coding in individual neurons (Extended Data Fig. 2c), we used the same basic GLM structure without value terms (*ν*_*j*_) and adjusted the goal-progress bins (*x*_*i*_) to reflect different condition groupings:

- Patch-level task-structure GLM: 24 patch ID × progression binary indicators
- Action-level task-structure GLM: 16 action × progression binary indicators
- Progression-level task-structure GLM: 8 goal-progress binary indicators

To test the significance of full models against null (intercept-only) models or no-value models, we used 10-fold cross validation and repeated 100 times per neuron. Additionally, L1 regularization was applied to prevent overfitting.

#### GLM comparison and neuron categorization (Extended Data Fig. 2b)

We determined the level of generality at which each neuron encoded switch value by pairwise comparison between GLMs fitted with (i) value × progression terms, (ii) value × progression × action terms, (iii) value × progression × patch ID terms, or (iv) value × progression × journey terms. All models included the same set of progression × journey indicators (i.e., the task-structure terms), ensuring that only the value-modulation structure varied across models. For each neuron, we compared the fraction of deviance explained by each pair of GLMs using one-sided Wilcoxon rank-sum test, with Holm– Bonferroni correction applied across comparisons. A neuron was assigned a value-coding generality category based on the following criteria:

- If one model explained significantly more fraction of deviance than all others, that model was selected.
- If multiple models each explained significantly more deviance than the remaining models but showed statistically indistinguishable performance, we selected the most parsimonious one among them.

This procedure yielded a principled categorization of each neuron according to the best-fitting and least complex model of its value-encoding generality.

#### Principal component analysis (PCA) and subspace definition

To explore population-level structure in mPFC activity of each day, we used PCA to identify neural dimensions capturing the largest variance within a specified analysis window (Extended Data Fig. 3). Specifically, neuronal firing rates were estimated by convolving spike trains of each single unit with a Gaussian kernel (σ = 50 ms). We then constructed a data matrix R (Conditions × Time) × Neurons, where conditions were defined by proximity to patch switches (Fig. 3a, c), proximity to patch switches × action (Fig. 3b, Extended Data Fig. 4c), proximity to patch switches × patch identity (Extended Data Fig. 4a, d), or proximity to patch switches × journey (Extended Data Fig. 4b, e). Firing rates of each neuron were averaged across all trials belonging to a given condition. The **navigation subspace** was identified by applying PCA to neural data from 0 to +3 s relative to trial initiation (i.e., the last nose-poke time at the starting port). This interval covered the median run period on stay trials. We then projected neural data from the same time window into this subspace (Fig. 3a, b; Extended Data Fig. 4a, b). To isolate dimensions more strongly linked to value-related variance, we performed a second PCA on neural activity from -0.8 to +0.2 s relative to trial initiation, an interval during which rats paused at reward ports on both rewarded and unrewarded trials before initiating the next movement (Extended Data Fig. 5). The leading PCs from this decomposition defined the **pre-move subspace**. We then projected neural data (i) during the -0.8 to +3 s window relative to trial initiation on stay trials and the -0.8 to +4.5 s window relative to trial initiation on switch trials into this subspace to reveal the continuous spiral trajectories across trials (Fig. 3c, Extended Data Fig. 4c-e), and (ii) during the -5 to 0 s interval from trial initiation after reward and the -0.8 to 0 s interval from trial initiation after omission to reveal evolution of neural trajectories spanning the outcome period of the preceding trial (Fig. 3d).

#### Latent Factor Analysis via Dynamical Systems (LFADS) for inferring single-trial neural dynamics

To identify single-trial neural trajectories in the latent state space, and particularly, the effect of individual reward outcomes on value updating, we applied LFADS to reconstruct denoised neural spiking data^55^. LFADS is a deep-learning-based sequential variational autoencoder that models neural population activity as noisy observations of an underlying latent dynamical system and produces smooth firing-rate estimates for each neuron. This approach facilitates analysis of single-trial population trajectories in neural state spaces.

We implemented LFADS using the AutoLFADS framework^56^, which trains multiple LFADS models in parallel on a single dataset and optimizes hyperparameters via population-based training^57^. At periodic evaluation steps, models were ranked according to performance on held-out validation data, and poorly performing runs were replaced by copies of top-performing runs with small stochastic perturbations to selected hyperparameters. We additionally used coordinated dropout to discourage trivial input passthrough solutions and to promote learning of shared population-level structure.

Each round of AutoLFADS was applied to a full day of neural recordings for a given animal, excluding rest sessions. Prior to model training, spike counts of each neuron were binned in 10 ms intervals. To reduce redundancy and mitigate overfitting, we excluded neurons whose spiking rates were highly correlated with others, such that pairwise cross-correlations of all retained neurons were below 0.1. Continuous recordings were segmented into 500 ms windows (i.e., 50 bins of spike counts) with 100 ms overlap. Every fifth segment was held out for validation, and the remainders were used for training. We then applied PCA to the LFADS-inferred firing rates to extract denoised, single-trial neural states (Fig. 3e).

#### Single-trial neural-state shifts after reward versus omission

LFADS-inferred firing rates of single trials were projected into a low-dimensional state space constructed by PCA (specifically, pre-move subspace in Fig. 3e). We focused on the two leading PC axes along which spiral neural trajectories across pre-switch and switch trials showed salient value-related gradients. Single-trial neural-state shifts were defined as the vector connecting neural states at initiation of two successive trials. We took the average of neural-state shifts after reward and omission separately and then calculated the cosine angle between these two mean vectors to measure their geometric relationship (Fig. 3f). To assess statistical significance, we bootstrapped single-trial neural-state shifts with replacement and recomputed the cosine angle between reward- and omission-evoked mean vectors for each resample.

#### Cross-validated switch value decoding from neural population activity

To quantify value representations in mPFC population activity, we decoded the *switch value* estimated by the Beta– Bernoulli behavioral model using LASSO regression with L1 regularization. Given the interaction between task structure and value in neuronal coding, we discretized each trial into multiple progression bins and averaged neuronal spiking data within each bin. Depending on the level of task abstraction (generality) examined, bins were defined as the follows.

- Progression-based value decoder (Fig. 4; Extended Data Fig. 7): all stay trials were pooled together and divided into 11 progression bins each (8 bins during run time and 3 bins during the outcome period).
- Action-based value decoder (Extended Data Fig. 7d): stay trials with the same actions (turning left or turning right) were grouped together and trials in each group were divided into 8 run-time progression bins (i.e., 16 progression × action bins total).
- Patch-based value decoder (Extended Data Fig. 7d): stay trials within the same patch were grouped together and trials in each group were divided into 8 run-time progression bins (i.e., 24 progression × patch bins total).
- Journey-based value decoder (Extended Data Fig. 7d): stay trials of the same journey type were grouped together and trials in each group were divided into 8 run-time progression bins (i.e., 48 progression × journey bins).

For each bin, we trained a decoder of the form:

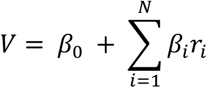

where *r*_*i*_ denotes the firing-rate vector of neuron *i* (out of N simultaneously recorded neurons) across all time points assigned to that bin, *β*_*i*_ is its decoding weight, and *V* is the corresponding vector of Beta-Bernoulli value estimates. All bins belonging to the same trial were assigned the same per-trial value estimate, so repeated entries in *V* reflect multiple time points from a single trial rather than moment-by-moment value updates. During decoder testing, only the first and last segments of switch trials were included and discretized into the same progression bins (see schematic in Fig. 4a).

To prevent overfitting, we implemented a leave-one-out cross-validation procedure: we grouped all trials by their proximity to a patch-switch decision (e.g., six visits away, five visits away, so on and so forth, from an upcoming switch) and used all trials except those in one of these groups to train the model; the held-out group (e.g., all trials that were two visits away from a switch) was used exclusively for testing. Critically, this procedure was repeated so that each stay-trial group had been left out once and all switch trials were always excluded from training. To compare decoding performance across decoders at different levels of generality (Extended Data Fig. 7d), we randomly subsampled trials when training the progression-based, action-based, and patch-based value decoders so that all models were fitted on matched numbers of trials. This ensured that differences in decoding accuracy reflected structure in the neural data rather than disparities in sample size.

#### Quantification of neural value repletion

To measure the degree to which neural value representations repleted when rats entered a new patch (Fig. 5b-d), we computed:

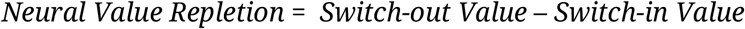

where *Switch-out Value* refers to the average decoded switch value across the four pre-choice bins right before rats switched out of a patch, and *Switch-in Value* is the average decoded switch value across the four post-choice bins when rats switched back to that same patch.

#### Quantification of neural value depletion

To measure the degree to which neural value representations depleted when rats received rewards (Fig. 5f-h), we computed:

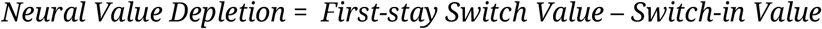

where *Switch-in Value* is the average decoded switch value across the four post-choice bins when rats switched to a new patch, and *First-stay Switch Value* refers to the average decoded switch value across the four post-choice bins of the subsequent first-stay trials in that same patch. We restricted analyses to cases where the switch-in trials were rewarded to focus on the effect of reward and to ensure comparability across early and late learning.

#### Quantification of neural value updating

To assess how reward outcomes influenced neural value representations, we defined two complementary metrics: (i) *Outcome Effect* on switch value was quantified as *Outcome Value* – *Final Switch-in Value*, where *Final Switch-in Value* is the decoded switch value in the final run-time bin (bin 8) of a switch trial, and *Outcome Value* denotes the mean decoded value in the outcome-period bins (during the pause time) of that same switch trial (Fig. 5i); (ii) *Neural Value Update* was quantified as *Initial First-stay Value* – *Final Switch-in Value*, where *Final Switch-in Value* is as defined in (i), and *Initial First-stay Value* denotes the decoded switch value in bin 1 of the subsequent stay trial (Fig. 5i). We then assessed the effect of reward outcomes on *Outcome Effect* and *Neural Value Update* by computing:

i. ΔOutcome effect = mean(Outcome effect|Reward=0) − mean(Outcome effect|Reward=1)
ii. ΔValue update = mean(Value update|Reward=0) − mean(Value update|Reward=1) which are shown in Fig. 5j.

#### Predicting choices from behavioral value estimates or neural value decoding

To quantify how well value variables predicted rats’ decisions, we fit logistic regression models (GLMs with a logit link function) to classify stay versus switch choices using either the switch value estimated by the behavioral Beta-Bernoulli model or the switch value decoded from neural population activity (Figs. 1m, 4d; Extended Data Fig. 7c). Models were fit separately for each animal. Each model used a single predictor—the behaviorally estimated or neurally decoded switch value of each trial—and output the probability of a switch decision. Model predictions were binarized using a decision threshold optimized to maximize overall prediction accuracy. To avoid biased evaluation of decoding accuracy due to unbalanced numbers of switch and stay trials, switch-decoding accuracy (sensitivity) and stay-decoding accuracy (specificity) were computed separately and then averaged (Fig. 4d, Extended Data Fig. 7c).

### Statistics

Statistical tests for each analysis are described in the corresponding Methods subsections and figure legends. Unless otherwise noted, all bootstrapping was done over 10,000 iterations to acquire 95% CIs. All model comparisons were performed on models trained with matched numbers of trials, achieved by random subsampling to ensure equivalent sample sizes. Statistical significance was assessed at α = 0.05, with significance levels indicated as: * P < 0.05, ** P < 0.01, and *** P < 0.001.

## Data and code availability

Upon publication, all data and code will be shared publicly via the Distributed Archives for Neurophysiology Data Integration (DANDI) Archive and GitHub repositories at https://github.com/LorenFrankLab. Further information necessary for analysis of this dataset is available upon request from the corresponding author(s).

## Acknowledgements

We thank A. Kiseleva, V. Perez, and M. Yusifova for administrative support, V. Kharazia for assistance with histology, L. Tong for assistance with spike curation, S. Gu for assistance with behavioral modeling, S. Tseng, D. J. O’Shea, S. Vyas, L. Tian, and D. Shin for analysis discussions, and A. K. Gillespie for rat illustrations. This work was supported by the Jane Coffin Childs Memorial Fund (X.S.), NSF GRFP 1650113 (A.E.C.), NIH F31MH124366 (A.E.C.), UCSF Jonas Cohler Discovery Fellowship (A.E.C.), the Life Sciences Research Foundation (A.J.), NIH F30MH126483 (J.A.G.), NIH R01MH121093 (N.D.D.), NIH R01MH136875 (J.D.B., N.D.D., L.M.F.), Simons Collaboration on the Global Brain 521921 and 542981 (L.M.F.), and Howard Hughes Medical Institute (L.M.F.).

## Author contributions

Conceptualization, X.S., A.E.C., N.D.D., L.M.F.; Methodology, X.S., A.E.C., A.E.K., T.A.K., J.D.B., N.D.D., L.M.F.; Data Collection, A.E.C., E.J.M.; Surgical Procedures, A.E.C., E.J.M., A.J., J.A.G.; Formal Analysis, X.S., A.E.C., A.E.K., C.B.W.; Software, X.S., A.E.C., A.E.K., E.J.M., E.L.D., N.D.D., L.M.F.; Resources, J.Z., P.T., J.H., A.Y., R.H., L.M.F.; Data Curation, X.S., A.E.C., E.J.M., L.M.F.; Writing – Original Draft, X.S., A.E.K., L.M.F.; Writing – Review & Editing, X.S., A.E.C., A.E.K., E.J.M., C.B.W., A.J., J.A.G., E.L.D., T.A.K., C.P., J.D.B., N.D.D., L.M.F.; Visualization, X.S., A.E.K., A.E.C.; Supervision, L.M.F.

## Declaration of interests

The authors declare no competing interests.

**Extended Data Fig. 1.**
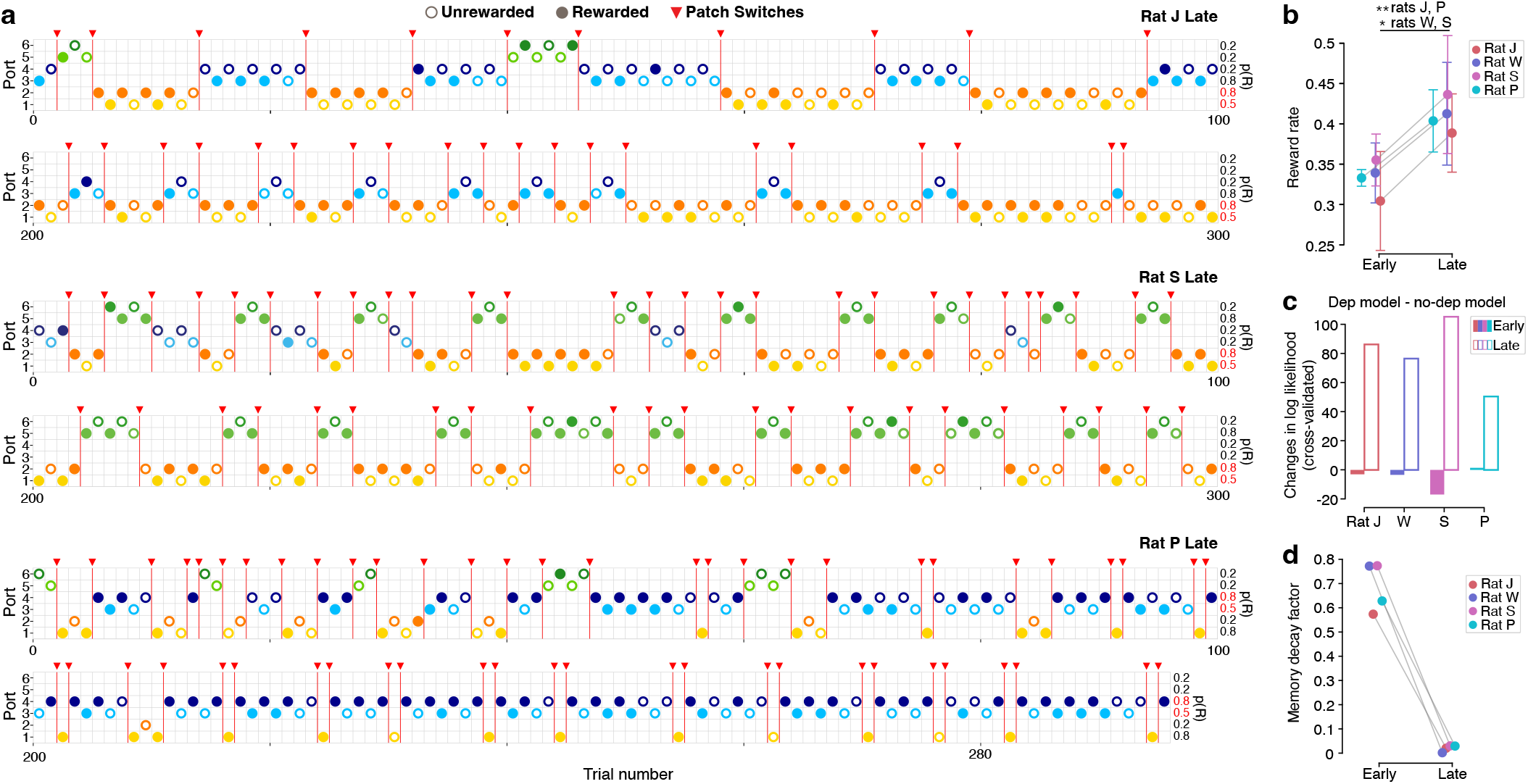
Behavioral performance of all rats and additional model metrics reflect the emergence of meta-learning. **a**, Example behavior from the first and last 100 trials of representative late-learning sessions in rats J, S, and P. Similar to rat W (Fig. 1c, bottom), these rats quickly identified the high-reward patch in late-learning sessions and alternated frequently between it and the medium-reward patch. The p(R) of the high-reward patch is marked in red. Filled circles: rewarded trials; hollow circles: unrewarded trials (colored by port identity for visualization). Red triangles: patch switches. **b**, As expected from meta-learning, reward rate increased significantly in late learning compared with early learning across all rats (one-sided Wilcoxon rank-sum test: P_J_ = 0.0098, P_W_ = 0.012, P_S_ = 0.016, P_P_ = 0.0013). **c**, Incorporating the depletion factor into the behavioral model improved its accuracy (i.e., higher cross-validated log-likelihood) in predicting decisions during late learning across all rats. **d**, The memory decay factor inferred from behavior decreased across days, reflecting improved retention of the underlying reward probability over learning.

**Extended Data Fig. 2.**
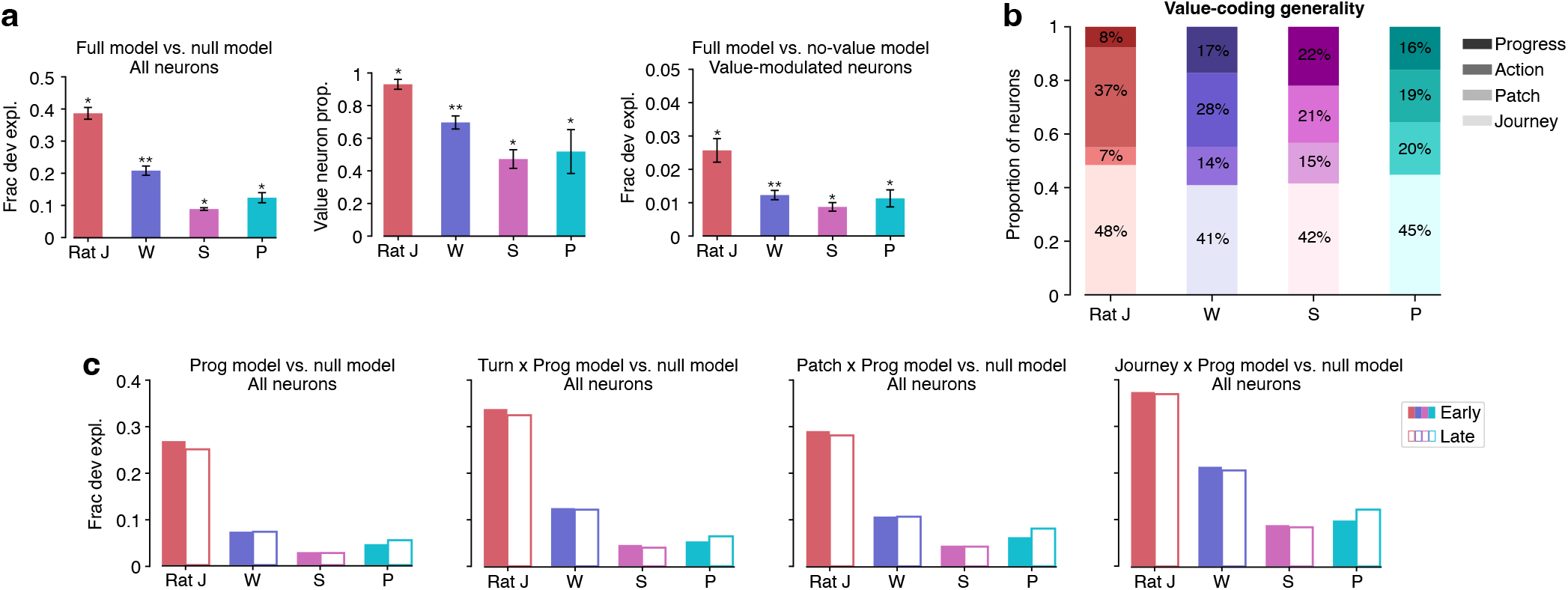
Quantification of GLM encoding patterns. **a**, GLM summary statistics across all days for each rat. Left: The full GLM (including value terms) explained a significantly greater fraction of deviance in single-neuron activity than the null (intercept-only) model (one-sided Wilcoxon signed-rank test > 0: P_J_ = 0.016, P_W_ = 0.0078, P_S_ = 0.016, P_P_ = 0.031). Middle: Proportion of value-encoding neurons (one-sided Wilcoxon signed-rank test > 0: P_J_ = 0.016, P_W_ = 0.0078, P_S_ = 0.016, P_P_ = 0.031). Right: The fraction of deviance attributable specifically to value terms in the full model was significantly greater than zero (one-sided Wilcoxon signed-rank test > 0: P_J_ = 0.016, P_W_ = 0.0078, P_S_ = 0.016, P_P_ = 0.031). Error bars: standard deviation. **b**, Categorization of value-modulated neurons by their value-coding generality. For each neuron, we performed pairwise comparisons among GLMs incorporating value × progression, value × progression × action, value × progression × patch ID, or value × progression × journey terms. The most parsimonious model whose fraction of deviance explained was not significantly lower than that of any competing model defined the neuron’s value-coding generality (one-sided Wilcoxon rank-sum test with Holm-Bonferroni correction; see Methods). **c**, Fraction of deviance explained by task-structure GLMs compared with the null model in early- and late-learning sessions. Models included goal-progress terms, turn × progression terms, patch ID × progression terms, and journey × progression terms, respectively, and excluded value terms. Deviance explained was averaged across neurons within each rat.

**Extended Data Fig. 3.**
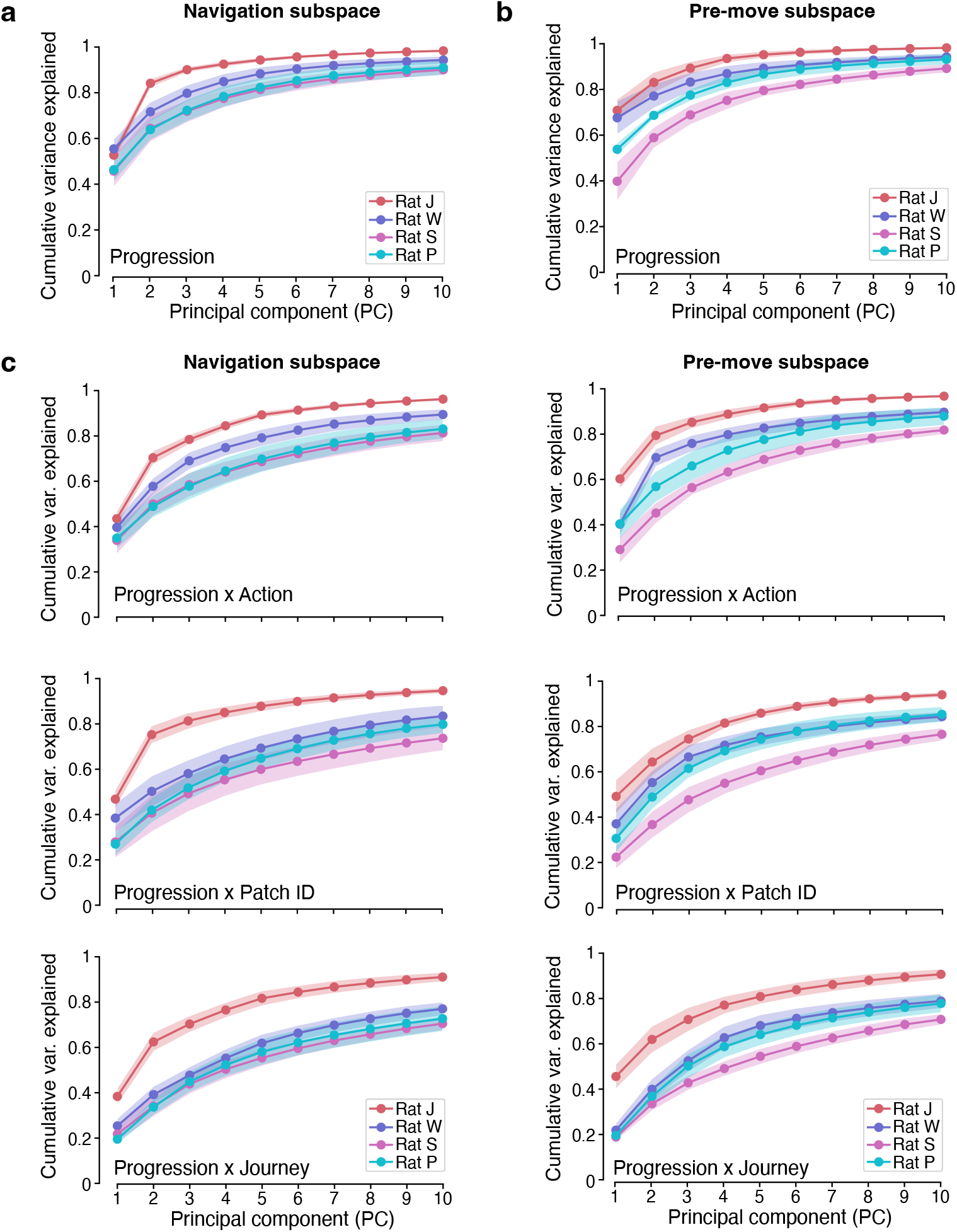
Variance explained by navigation and pre-move neural subspaces constructed by PCA. **a**, Cumulative variance explained by the leading PCs of run-time neural data grouped by proximity to a switch trial. **b**, Cumulative variance explained by the leading PCs of pre-move neural data (the end of pause time before movement initiation) grouped by proximity to a switch trial. **c**, Cumulative variance explained by the leading PCs of run-time and pre-move neural data grouped by different task conditions, including action × switch proximity (top), patch ID × switch proximity (middle), and journey × switch proximity (bottom). **a-c**, Shaded areas: standard deviation across all days for each rat.

**Extended Data Fig. 4.**
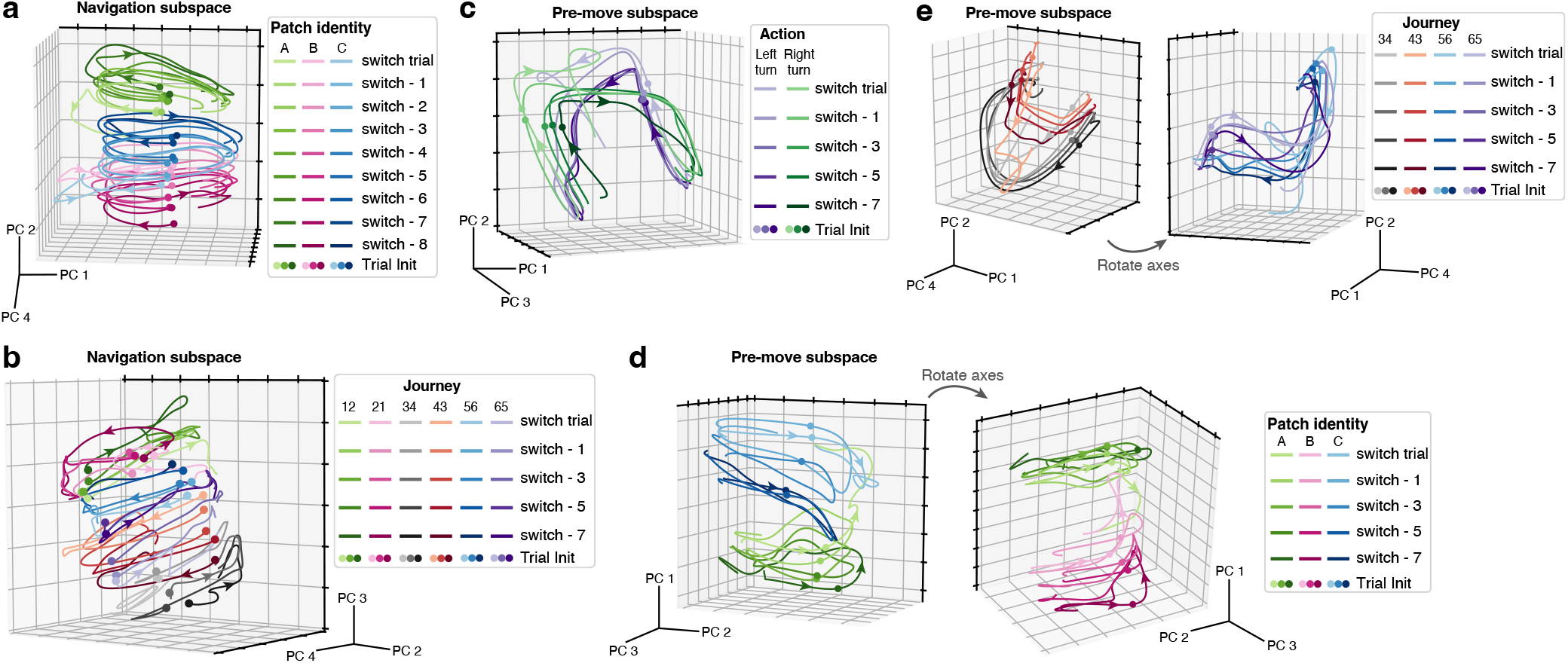
Generalized dynamical motifs across task conditions. **a**, In the navigation subspace, progress-tracking trajectories for each foraging patch were orderly displaced according to proximity to a switch, which also correlated with the switch value gradient. **b**, Progress-tracking trajectories similarly sorted by switch proximity for each individual journey in the navigation subspace. Every other pre-switch trial is shown for clearer visualization. **c**, In the pre-move subspace, neural trajectories expressed a value × progression × action motif, with value-related displacement across pre-switch and switch trials. Every other pre-switch trial is shown for clearer visualization. **d**, Dynamical motifs tracking value × progression × patch ID in the pre-move subspace. Showing every other pre-switch trial and two patches at a time for clearer visualization. **e**, Dynamical motifs tracking value × progression × journey in the pre-move subspace. Showing every other pre-switch trial and two pairs of journeys for clearer visualization. **a-e**, All sessions from one example day are shown. Circles: trial initiation. Arrowheads: direction of neural trajectory evolution.

**Extended Data Fig. 5.**
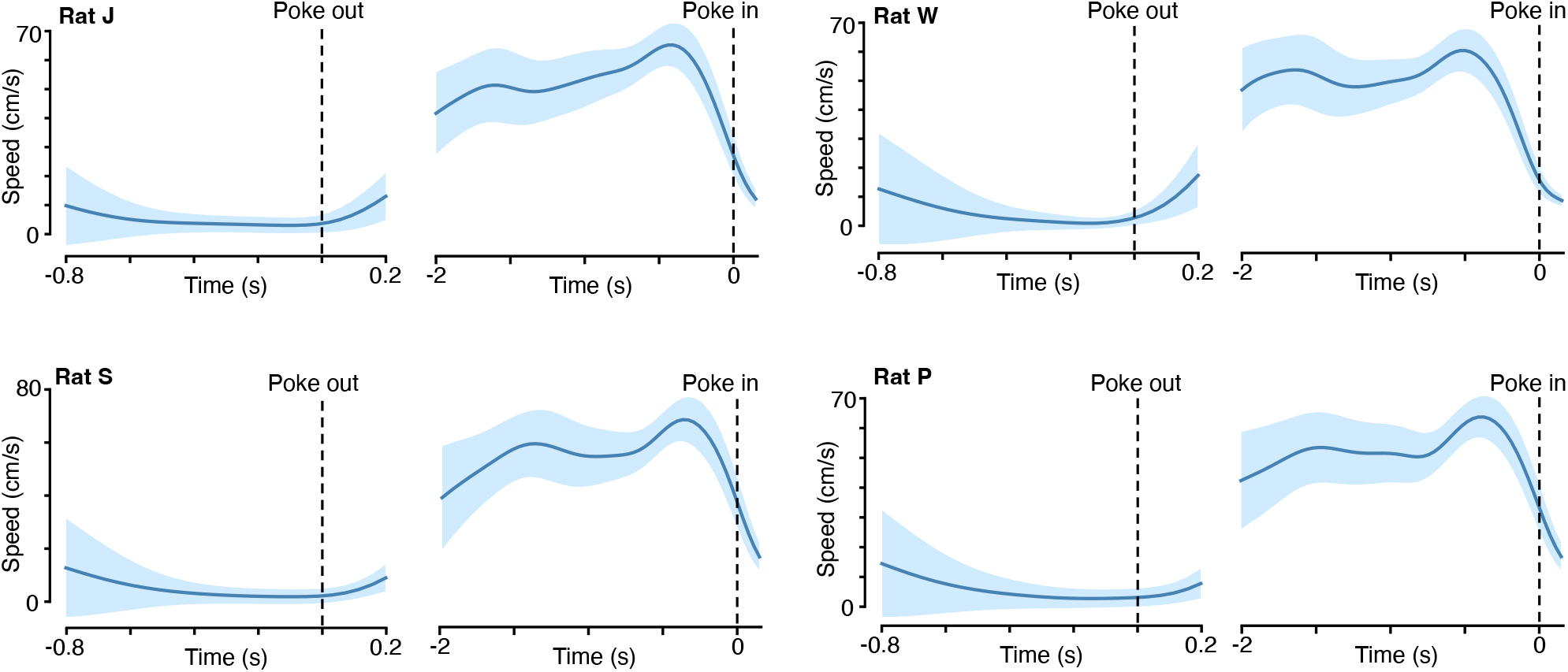
Rats’ movement speed during pre-move and run periods. Left panel for each rat: Mean speed remained at or below ∼10 cm/s throughout the pre-move time window used to construct the pre-move neural subspace, indicating minimal locomotion. Such immobility during this period allowed extraction of neural variance primarily related to value. ‘Poke out’ marks the time when rats withdrew from the reward port. Right panel for each rat: During the run period of each trial, mean speed remained above ∼20 cm/s until rats approached the reward port, consistent with sustained locomotion throughout this interval. ‘Poke in’ marks the moment rats arrived at the port. Shaded areas: standard deviation across all days for each rat.

**Extended Data Fig. 6.**
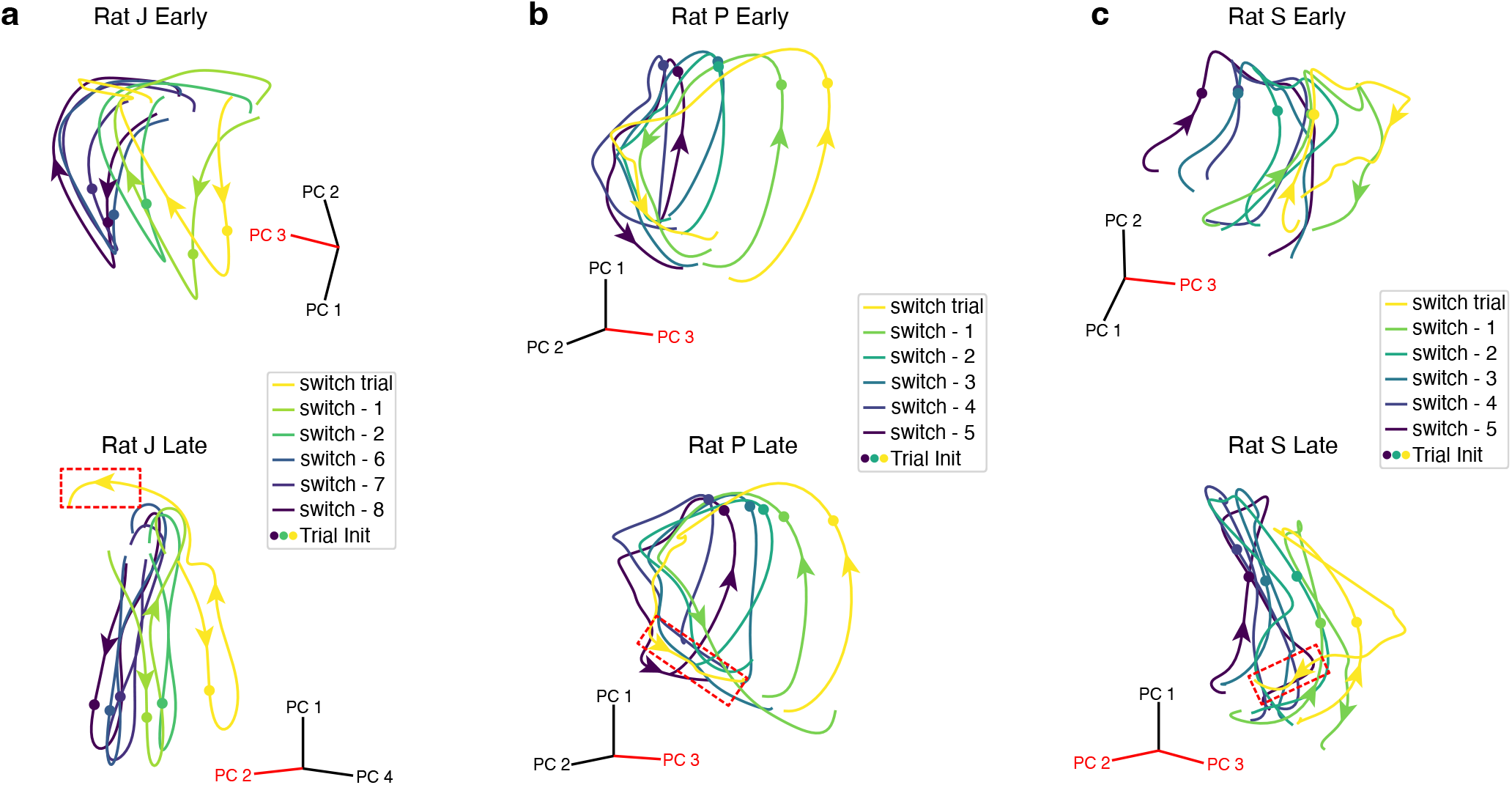
Group-averaged neural trajectories in the pre-move subspace during early and late learning. Average neural trajectories of trials grouped by proximity to a switch decision. For each rat, two example days are shown. The PC axes along which we observed prominent value-related displacement of spiral trajectories are highlighted in red. The resetting trajectories corresponding to entry into the final segment of a new patch (before reward outcome delivery) are emphasized within rectangular boxes. In early-learning sessions, rats J and S showed minimal resetting upon entering a new patch, whereas rats W (see Fig. 3c) and P exhibited partial resetting that was weaker than that observed in late-learning sessions. Note that although PCA is useful for visualizing dynamical structure, variability in trial counts and the number of recorded neurons can affect the apparent geometry of low-dimensional trajectories (e.g., occasional entanglement of spiral gradients in late sessions for rats J and S). Importantly, our conclusions primarily relied on decoding-based representational analyses, which provided a more robust quantification of value-coding structure. Nevertheless, PC-space trajectories remain informative for interpreting the organization and evolution of the underlying neural dynamics.

**Extended Data Fig. 7.**
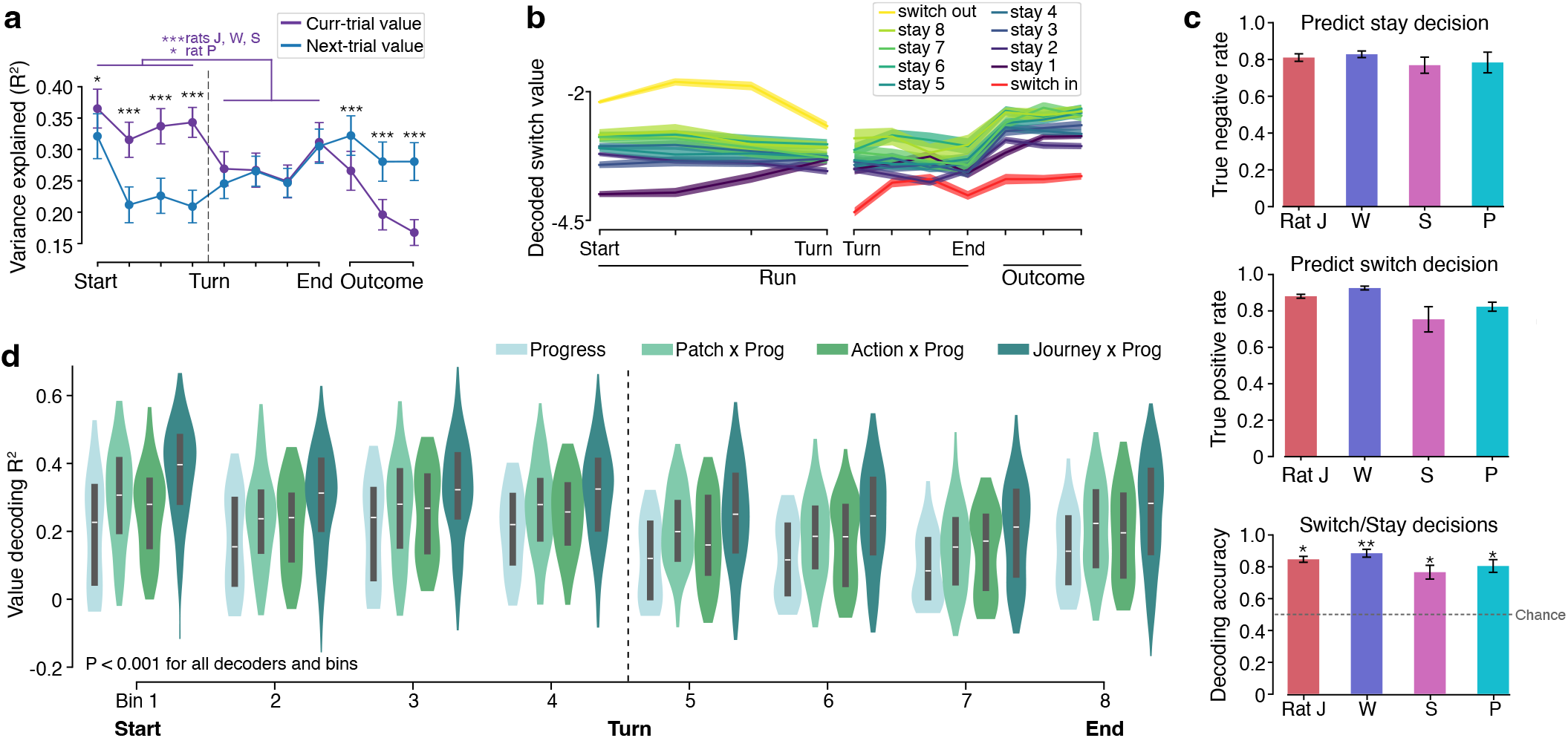
Additional measurements of switch value decoding. **a**, Variance of switch value (estimated by the behavioral model) explained by the neural decoder in each of the 11 progression bins. Data from all sessions and rats were pooled for statistical testing. In pre-choice bins (1-4), the decoder captured significantly more variance of the *current-trial value* than of the *next-trial value* (paired one-sided Wilcoxon signed-rank test: P_bin1_ = 0.011, P_bin2_ = 6.35 × 10^-5^, P_bin3_ = 3.28 × 10^-6^, P_bin4_ = 5.96 × 10^-8^). The relationship reversed in outcome-period bins (9-11), with the decoder explaining more variance of the next-trial value (paired one-sided Wilcoxon signed-rank test: P_bin9_ = 4.17 × 10^-6^, P_bin10_ = 1.19 × 10^-7^, P_bin11_ = 5.96 × 10^-8^). When decoding the current-trial value (purple), variance explained was higher in pre-choice than in post-choice bins (permutation test per animal: P_J_ = 0.0, P_W_ = 0.0, P_S_ = 0.0006, P_P_ = 0.0248). **b**, Decoded switch value across 11 progression bins for stay and switch trials on an example day (5 sessions). Value during the outcome period appeared to link the terminal value of one trial to the initial value of the next, consistent with value updating. Shaded areas: SEM across trials. **c**, Decoding accuracy for stay decisions (top), switch decisions (middle), and the mean of both (bottom) across all days for each rat. Accuracy remained stable across days. One-sided Wilcoxon signed-rank test > 0.5 (bottom): P_J_ = 0.016, P_W_ = 0.0078, P_S_ = 0.016, P_P_ = 0.031. Error bars: standard deviation. **d**, Variance of switch value explained by neural decoders trained at different levels of task generality (during run time): 8 progression bins, 16 action × progression bins, 24 patch × progression bins, and 48 journey × progression bins. We subsampled trials before fitting the progression-, action-, and patch-based decoders to ensure that the performance of all decoders was comparable with matched training and test data sizes. Variance of switch value explained by all decoders was significantly greater than zero (all days of all rats pooled for statistical testing; one-sided Wilcoxon signed-rank test: P < 0.001 across all progression bins, action × progression bins, patch × progression bins, and journey × progression bins).

**Extended Data Fig. 8.**
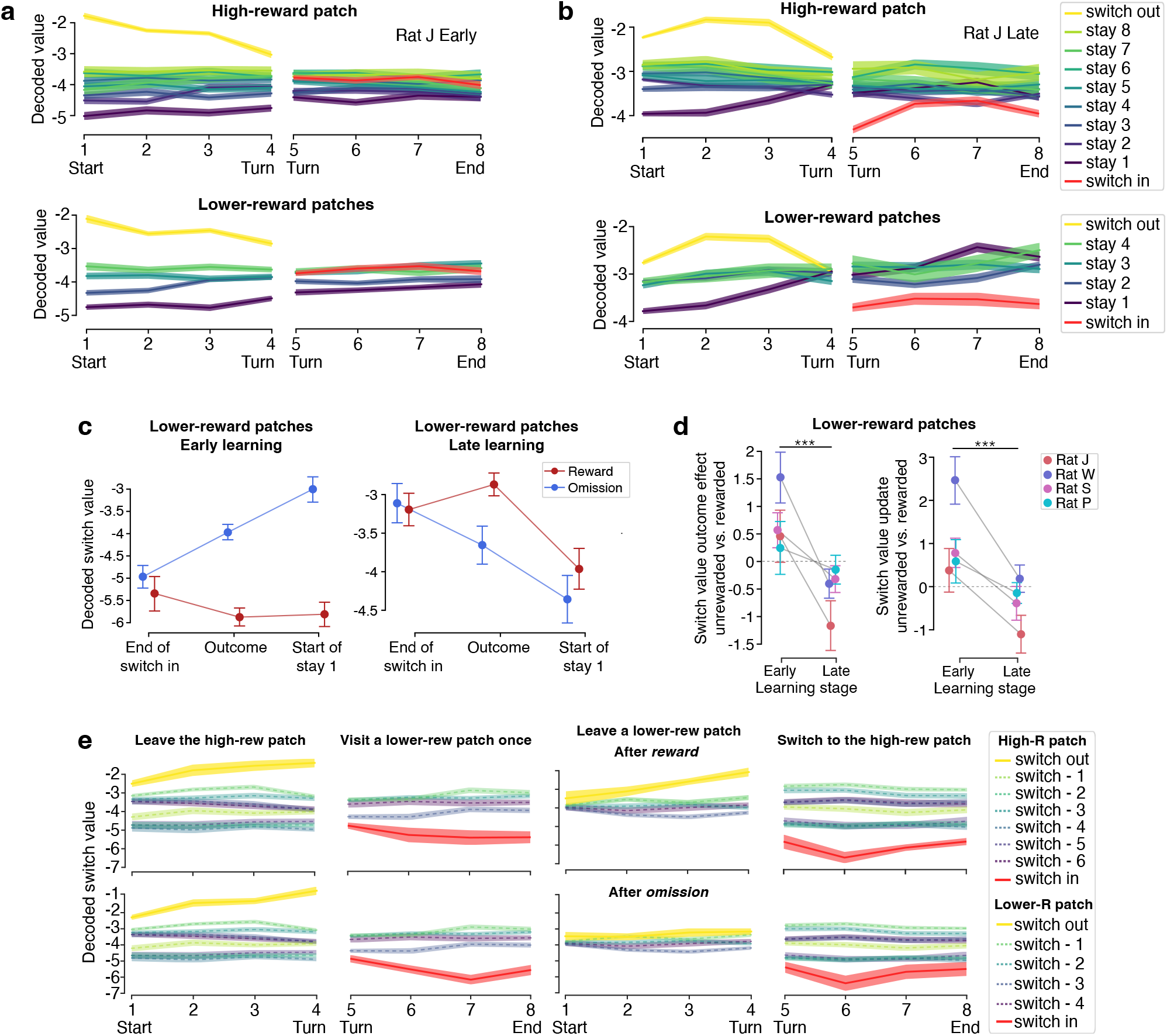
Changes in neural value representations at finer timescales and in lower-reward patches are consistent with the meta-learned strategy. **a, b**, Decoded switch value across run-time progression bins within a patch for switch-in (red), the subsequent stay (violet to green), and switch-out (yellow) trials. Shown are early- and late-learning sessions from an example rat. Trials in the high-reward and lower-reward patches were analyzed separately because their average bout lengths typically differed (Fig. 1i). In the high-reward patch, decoded switch value during switch-in trials reset more substantially in late learning than in early learning (signature of inferring repletion). In both high-reward and lower-reward patches, decoded switch value increased from switch-in to first-stay trials in late learning (signature of anticipating depletion). Shaded areas: SEM across trials. **c, d**, Same measurements as in Fig. 5i, j for lower-reward patches. **d**, Linear mixed effects model testing random slopes per animal: outcome-effect difference—P_J_ = 9.05 × 10^-6^, P_W_ = 4.97 × 10^-12^, P_S_ = 6.78 × 10^-4^, P_P_ = 0.18; value-update difference— P_J_ = 4.35 × 10^-5^, P_W_ = 5.78 × 10^-12^, P_S_ = 3.19 × 10^-4^, P_P_ = 0.0075. Fixed slope across animals: outcome-effect difference, P = 3.80 × 10^-14^; value-update difference, P = 3.13 × 10^-14^. Error bars: 95% CI. **e**, Decoded switch value across run-time progression bins during single-visit episodes (solid curves) on an example day. For comparison, decoded switch value during stay bouts (i.e., when rats stayed for more than a single visit) in the same patch is shown as washed-out dashed curves. After both rewarded and unrewarded single visits, decoded switch value at the start of the subsequent trial was, on average, higher than that during stay bouts (third column). Shaded areas: SEM across trials.

**Extended Data Fig. 9.**
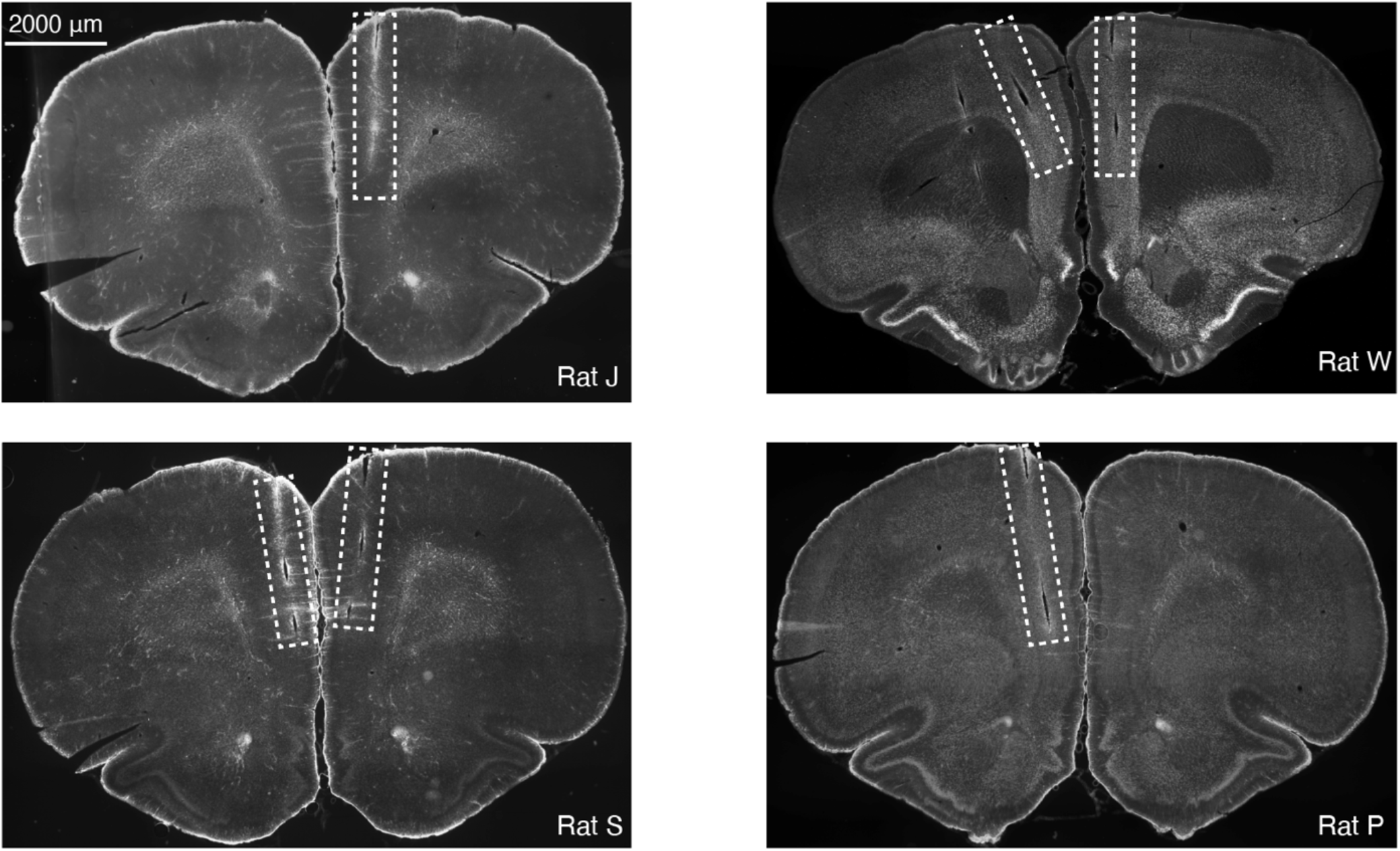
Implant locations. Example PFC histological sections from rats J, W, S, and P. Tracks of the mPFC probes are emphasized within rectangular boxes.

